# Novel Paju Apodemus Paramyxovirus 1 and 2, Harbored by *Apodemus agrarius* in The Republic of Korea

**DOI:** 10.1101/2021.03.03.433816

**Authors:** Seung-Ho Lee, Jin Sun No, Kijin Kim, Shailesh Budhathoki, Kyungmin Park, Geum Young Lee, Seungchan Cho, Hyeok Sun Choi, Bong-Hyun Kim, Seunghee Cho, Jong Woo Kim, Jin Gyeong Lee, Seung Hye Cho, Heung-Chul Kim, Terry A. Klein, Chang-Sub Uhm, Won-Keun Kim, Jin-Won Song

## Abstract

Paramyxoviruses, negative-sense single-stranded RNA viruses, pose a potential threat to public health. Currently, 78 species and 17 genera of paramyxoviruses are classified and harbored by multiple natural reservoirs, including rodents, bats, birds, reptiles, and fish. *Jeilongvirus* has been proposed as a novel paramyxovirus genus containing J-, Beilong, and Tailam viruses, found in wild rodents. Using RT-PCR, 824 *Apodemus agrarius* individuals were examined for the prevalence of paramyxovirus infections. Paramyxovirus RNA was detected in 108 (13.1%) rodents captured at 14 trapping sites in Korea. We first present two genetically distinct novel paramyxoviruses (genus *Jeilongvirus*), Paju Apodemus paramyxoviruses 1 (PAPV-1) and 2 (PAPV-2), from *A. agrarius*. Six PAPV strains were completely sequenced using next-generation and Sanger sequencing. PAPV-1 genome comprised 19,716 nucleotides, with eight genes (3′-N-P/V/C-M-F-SH-TM-G-L-5′), whereas PAPV-2 genome contained 17,475 nucleotides, with seven genes (3′-N-P/V/C-M-F-TM-G-L-5′). The disparity between PAPV-1 and -2 revealed the presence of the *SH* gene and length of the *G* gene in the genome organization. The phylogenies of PAPV-1 and -2 belong to distinct genetic lineages of *Jeilongvirus* despite being from the same natural host. PAPV-1 clustered with Beilong and Tailam viruses, while PAPV-2 formed a genetic lineage with Mount Mabu Lophuromys virus-1. PAPV-1 infected human epithelial and endothelial cells, facilitating the induction of type I/III interferons, interferon-stimulated genes, and proinflammatory cytokines. Therefore, this study provides profound insights into the molecular epidemiology, virus-host interactions, and zoonotic potential of novel rodent-borne paramyxoviruses.

**Importance:** Paramyxoviruses are a critical public health and socio-economic burden to humans. Rodents play a crucial role in transmitting pathogens to humans. In the last decade, novel paramyxoviruses have been discovered in different rodents. Here, we found that *Apodemus agrarius* harbored two distinct genotypes of the novel paramyxoviruses, Paju Apodemus paramyxovirues 1 (PAPV-1) and 2 (PAPV-2), possessing unique genome structures that are responsible for encoding TM and G proteins of different sizes. In addition, PAPV-1 infected human epithelial and endothelial cells, facilitating the induction of type I/III IFNs, ISGs, and proinflammatory cytokines. Thus, this study provides significant insights into molecular prevalence, virus-host interactions of paramyxoviruses. These observations raise the awareness of physicians and scientists about the emergence of new rodent-borne paramyxoviruses.

## Introduction

Zoonotic diseases, transmitted from reservoir hosts to humans, comprise the majority of emerging and re-emerging infectious diseases and are public health and socio-economic threats (1–3). Emerging outbreaks of zoonotic viruses, such as severe acute respiratory syndrome coronavirus 2, have increased recently because of expanding human activities that have enabled virus spillover, particularly in situations that facilitate close contact among diverse wildlife species, domesticated animals, and humans (4). Rodents serve as potential mammalian hosts and pose the highest risk of harboring zoonotic viruses to date (3). These animals cause significant economic losses in agriculture and transmit infectious agents including viruses, bacteria, and parasites that cause hemorrhagic fever, tsutsugamushi disease, and leptospirosis (5, 6). Among the rodents in Asia and Europe, *Apodemus* species is a natural reservoir host carrying pathogens that are detrimental to humans, and *A. agrarius* is widely distributed in various natural environments (e.g., rural areas, agricultural fields, and forests). Metagenomic studies and continuous surveillance of potential viruses in small mammals provided clues for preventive and mitigative strategies against new emerging and re-emerging infectious diseases (7–12).

Paramyxoviruses are non-segmented, negative-sense single-stranded RNA viruses. *Paramyxoviridae* is divided into four subfamilies: *Avulavirinae*, *Rubularvirinae*, *Metaparamyxovirinae*, and *Orthoparamyxovirinae*. *Orthoparamyxovirinae* is classified into nine genera: *Respirovirus*, *Aquaparamyxovirus*, *Fetavirus*, *Henipavirus*, *Jeilongvirus*, *Narmovirus*, *Salemvirus*, *Sunshinevirus*, and *Morbillivirus* (13). Paramyxoviruses have a wide host range, including vertebrates (mammals, birds, reptiles, and fish) (14). Some paramyxoviruses, for example, human parainfluenza, Hendra, Nipah (NiV), mumps, and measles viruses, pose critical public health and socio-economic burdens owing to their pathogenicity in humans.

The newly established genus *Jeilongvirus* has been identified in all rodents, and it consists of seven recognized species, J-virus (JV) (15), Beilong virus (BeiV) (16), Tailam virus (TaiV) (17), Mount Mabu Lophuromys virus 1 (MMLV-1) and 2 (MMLV-2), Shaan virus, and Pohorje Myodes paramyxovirus 1 (PMPV-1) (18). In 1972, JV was isolated from the kidney autoculture of a moribund house mouse (*Mus musculus*) captured in northern Queensland, Australia (19). BeiV was first identified in a human mesangial cell line (16). In 2012, the presence of BeiV was confirmed in the kidney and spleen tissues of brown (*Rattus norvegicus*) and black (*Rattus rattus*) rats in Hong Kong (20). TaiV was isolated from the kidney and spleen tissues of Sikkim rats (*Rattus andamanensis*) at the Tai Lam country park in Hong Kong (17). MMLV-1 and -2 were discovered in the kidney of a Rungwe brush-furred rat (*Lophuromys machangui*) in Mozambique, and PMPV-1 was found in the kidney of a bank vole (*Myodes glareolus*) in Slovenia (18). These viruses possess two additional transcription units encoding the small hydrophobic (SH) and transmembrane (TM) proteins between the fusion (F) and receptor binding protein genes, with the exception of MMLV-1 and -2, which contain the *TM* gene but not the *SH* gene. In a previous study, paramyxovirus SH proteins were found to modulate the *in vitro* expressions of proinflammatory cytokines interleukin 6 (IL-6) and IL-8 via nuclear factor-κB (NF-κβ) activation (21). In particular, the SH protein of JV inhibited tumor necrosis factor-α (TNF-α) production and apoptosis *in vitro* and *in vivo* (22–24). However, the pathogenicity of these Jeilongviruses remains unexplored in humans.

In this study, 824 *A. agrarius* individuals were collected at 14 trapping sites and investigated for the prevalence, phylogenetic diversity, and genomic characterization of novel Paju Apodemus paramyxoviruses (PAPVs), PAPV-1 and -2, in the Republic of Korea (ROK). PAPV-1 infected human cells and induced the expression of type I/III interferons (IFNs), interferon-stimulated genes (ISGs), and proinflammatory cytokines. Thus, this study provides significant insights into the genetic diversity, evolutionary dynamics, and virus-host interactions of novel rodent-borne paramyxoviruses.

## Results

### Molecular screening and isolation of PAPVs in rodents

A total of 824 *A. agrarius* individuals were captured in various regions of the ROK from 2016 to 2018 **(Figure S1)**. Paramyxovirus RNA was detected in 81/824 (9.8%) *A. agrarius* individuals using specific primers targeting the genera *Respirovirus*, *Morbillivirus*, and *Henipavirus* (14,287–14,725 nt). Viral RNA was detected in 59/824 (7.2%) rodents via RT-PCR by targeting pan-*Orthoparamyxovirinae* (15,369–15,898 nt) (**Table S1)**. In total, 108 (13.1%) *A. agrarius* individuals were positive for paramyxovirus **(Table 1)**. The prevalence of PAPV-1 was 87/824 (10.6%), whereas that of PAPV-2 was 21/824 (2.6%) (*P* <0.001 in Fisher’s exact test). The geographic prevalence of PAPV was as follows: 39/361 (10.8%) in Gangwon Province, 63/434 (14.5%) in Gyeonggi Province, 1/6 (16.7%) in Gyeongsangnam Province, and 5/23 (21.7%) in Chungcheongnam Province **(Table S2)**. The rodent-borne paramyxovirus, PAPV, was first identified in *A. agrarius* captured in Paju, ROK. Among 370 male and 453 female *A. agrarius*, 56 (15.1%) and 52 (11.5%) harbored PAPV, respectively, indicating the gender-specific prevalence of this virus. Adult and old *A. agrarius* (20–30 g and 31–40 g, respectively) showed PAPV infection rates of 18.1% (45 animals) and 22.4% (34 animals), respectively, whereas the juvenile and subadult animals (˂20.0 g) showed low PAPV infection rates (6.7% and 4.5%, respectively). PAPV-positive rodents were found in spring, summer, autumn, and winter.

**Table 1.**
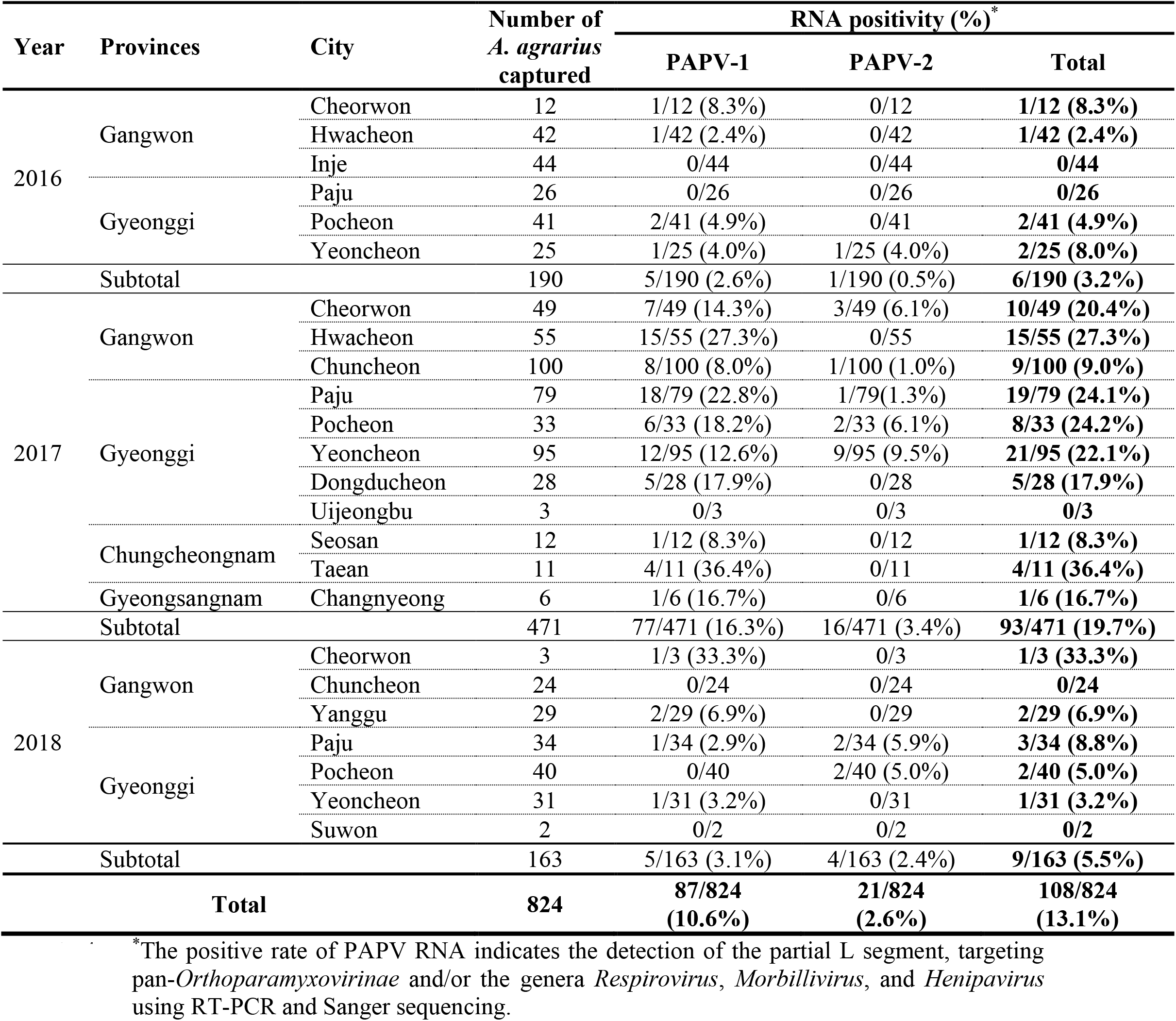
Prevalence of paramyxovirus infection based on Paju Apodemus paramyxoviruses (PAPVs) captured from 2016 to 2018 in the Republic of Korea.

*A. agrarius*-borne paramyxovirus was isolated from the kidney tissues of Aa17-179 and Aa17-297 using a cell culture-based method. The paramyxovirus isolates from the infected rodents showed a cytopathic effect of syncytia formation (not shown) in Vero E6 cells. The first isolate of PAPV-1 was confirmed by passaging two times for 14 days post-inoculation. The particles of PAPV-1 were observed using a transmission electron microscope **(Figure 1A)**. In addition, the number of infectious PAPV particles was 3×10^5^ PFU/mL, quantified using the plaque assay **(Figure 1B)**.

**Figure 1.**
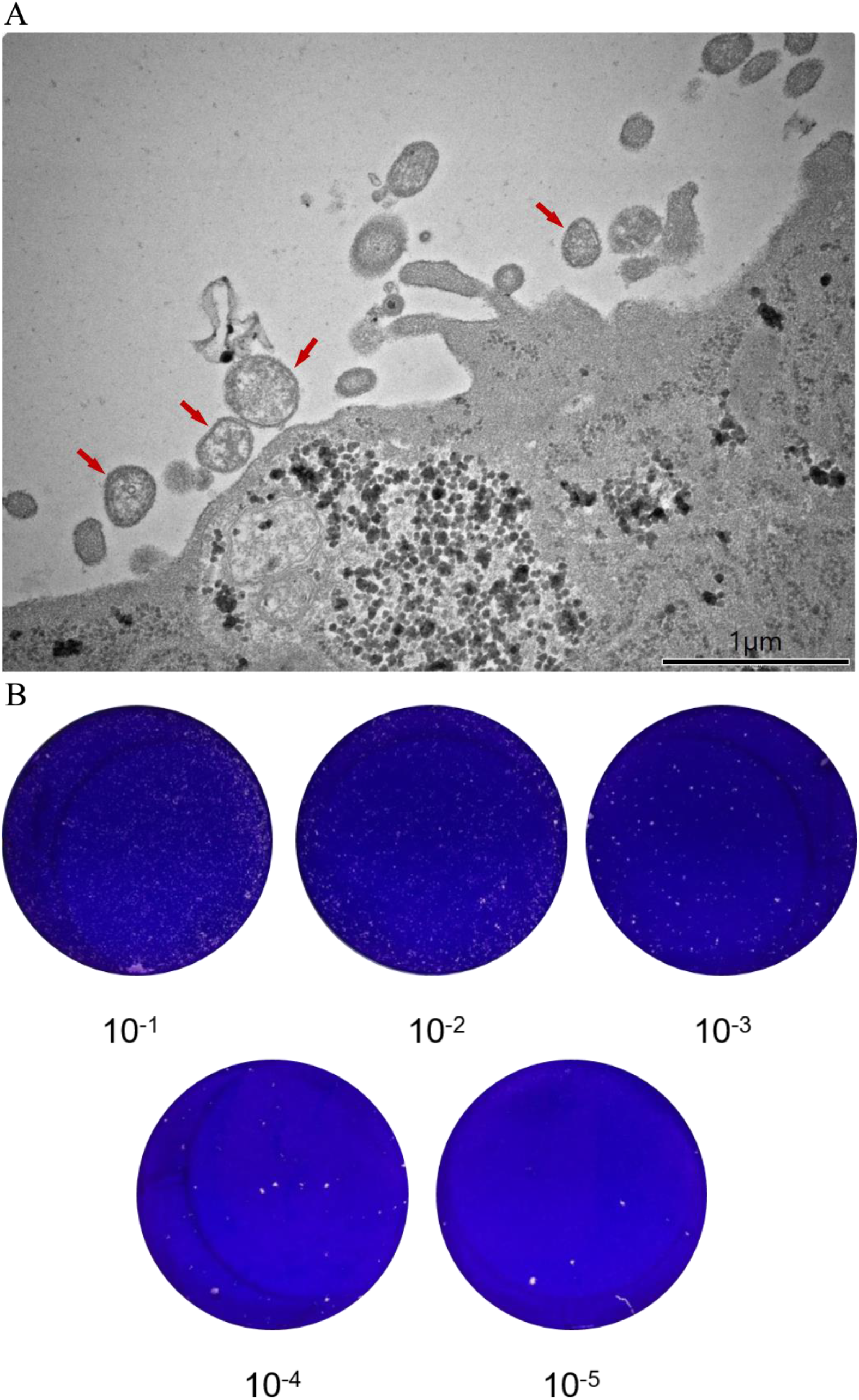
An electron microscopic image and the plaque assay of Paju Apodemus paramyxovirus 1 (PAPV-1) (A) PAPV-1 was imaged using transmission electron microscopy (TEM). (B) Photograph of a representative plaque assay plate of PAPV-1 inoculated into Vero E6 cells at 5 days post-infection. This single plate represents dilutions (from top-left to right: undiluted and dilutions at 1:10^1^, 1:10^2^, and bottom-left to right: dilutions at 1:10^3^, 1:10^4^, 1:10^5^) of the virus that mostly destroys the cell monolayer, producing the appropriate number of plaques to count.

### Whole-genome sequencing of PAPVs using next generation sequencing (NGS) and rapid amplification of cDNA ends (RACE) PCR

To obtain whole-genome sequences of PAPV, sequence-independent, single-primer amplification-based MiSeq of the Aa17-179 and Aa17-297 isolates generated eight contigs (520–976 nt in length) with significant similarities to the genomic sequence of paramyxoviruses. The NGS of Aa17-179 and Aa17-297 generated 1,623,052 and 1,479,714 viral reads, respectively, and the depth of the viral genome sequence was 144,317 and 79,344, respectively **(Table S3)**. The nearly complete genome sequences of four PAPV strains (Aa17-255, Aa17-260, Aa17-154, and Aa17-166) were acquired via Illumina sequencing. Both the 3′ and 5′end sequences of the viral genomes revealed incomplete complementary sequences with differences at nucleotide residues 4, 5, and 12. The genomic sequences of PAPV-1 and -2 have been deposited in GenBank (Accession number: MT823459-MT823464).

### Genomic organization of PAPVs

The whole genomes of PAPV-1 and -2 were 19,716 and 17,475 nt in length, with GC contents of 39.96–40.09% and 37.34%, respectively. PAPV-1 contained a genome structure composed of eight genes in the order of 3′-N-P/V/C-M-F-SH-TM-G-L-5′, while the genome structure of PAPV-2 comprised seven genes in the order of 3′-N-P/V/C-M-F-TM-G-L-5′ **(Figure 2)**. The N, M, F, G, and L genes encode one protein, while the P gene, in addition to the viral phosphoprotein, encodes some accessory proteins that arise through leaky scanning (C protein) or RNA editing (V/W protein). This RNA editing occurs through the addition of one or more guanine residues during transcription, following the recognition of a conserved RNA editing site. PAPV-1 and -2 were found to possess a putative RNA editing site (TTAAAAAAGGCA) within their P gene. This sequence matched a conserved motif sequence (YTAAAARRGGCA) found in all members of the genera *Henipavirus* and *Morbillivirus*, as well as in JV, TaiV, BeiV, and other rodent paramyxoviruses. PAPV-1 showed additional open reading frames between the F and G genes, encoding an SH and/or TM protein. In contrast, PAPV-2 showed the *TM* gene but not the *SH* gene. The 3′ leader and 5′ trailer sequences were 55 and 28 nt in length, respectively. The gene start, stop, and intergenic region sequences of PAPVs are shown in the Table S4.

**Figure 2.**
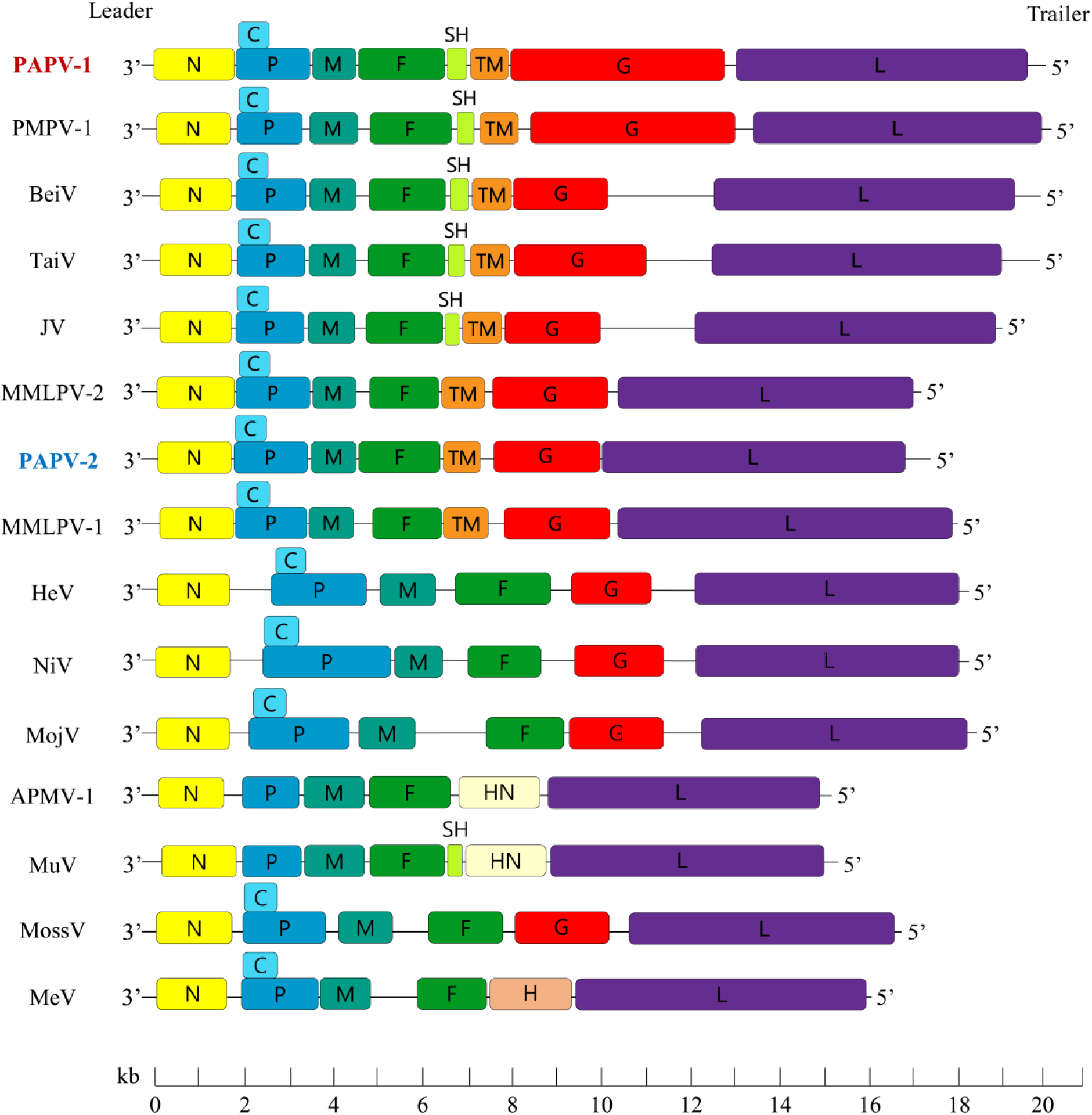
Organization of the genomes of Paju Apodemus paramyxoviruses 1 and 2. The genomic configurations of different paramyxoviruses are shown. The genome of paramyxovirus constitutes 8 to 9 coding regions, 3′ NP-C-P-M-F-SH-TM-G-HN-H-L 5′. The color boxes represent coding regions for each gene; N, yellow; C, sky blue; P, blue; M, viridian; F, green; SH, yellow green; TM, orange; G, red; HN, light yellow; H, Chilean pink, and L, purple. The genome size scale is provided at the bottom. Adobe Illustrator CS6 (http://www.adobe.com/products/illustrator.html) was used to construct the figures.

### Phylogenetic analysis of the novel PAPV strains

Phylogenetic inference of the whole-genome sequences of PAPVs demonstrated two distinct genotypes within Jeilongviruses **(Figure 3)**. The genetic cluster of PAPV-1 showed a high similarity (63.7–63.8%) with TaiV, while the PAPV-2 group shared a common ancestor with MMLV-1, with a genomic similarity of 71.6% **(Table S5)**. In addition, the amino acid sequences of the individual coding proteins of PAPV-1 and -2 constituted comparable phylogenetic patterns with the viral RNA genome sequences **(Figure S2)**. The partial L genomic sequences (14,287–14,725 nt) of novel PAPV strains phylogenetically belong to the genus *Jeilongvirus*, subfamily *Orthoparamyxovirinae*. Consistently, the phylogenies of PAPV formed two distinct genotypes compared to other Jeilongviruses. The PAPV-1 strains were closely related to TaiV, BeiV, and PMPV-1, whereas the PAPV-2 strains showed an independent genetic clustering with MMLV-1. The partial L gene sequences of PAPV-1 were differentiated into four genetic lineages **(Figure S3)**. The genetic lineage I of PAPV-1 originated geographically in Dongducheon, Paju, Pocheon, and Yeoncheon in Gyeonggi Province and Cheorwon in Gangwon Province. The genetic lineage II contained PAPV-1 strains in Pocheon in Gyeonggi Province, Chuncheon in Gangwon Province, and Taean and Seosan in Chungcheongnam Province. Additionally, genetic lineage III was found in Dongducheon and Paju in Gyeonggi Province, and in Hwacheon and Yanggu in Gangwon Province. A distinct strain from Changnyeong in Gyeongsangnam Province belonged to the genetic lineage IV. Further, the partial L gene sequences of PAPV-2 showed two phylogenetic clusters. Genetic lineage I included PAPV-2 in Yeoncheon, Paju, and Dongducheon, while genetic lineage II of PAPV-2 was observed in only Yeoncheon, Gyeonggi Province.

**Figure 3.**
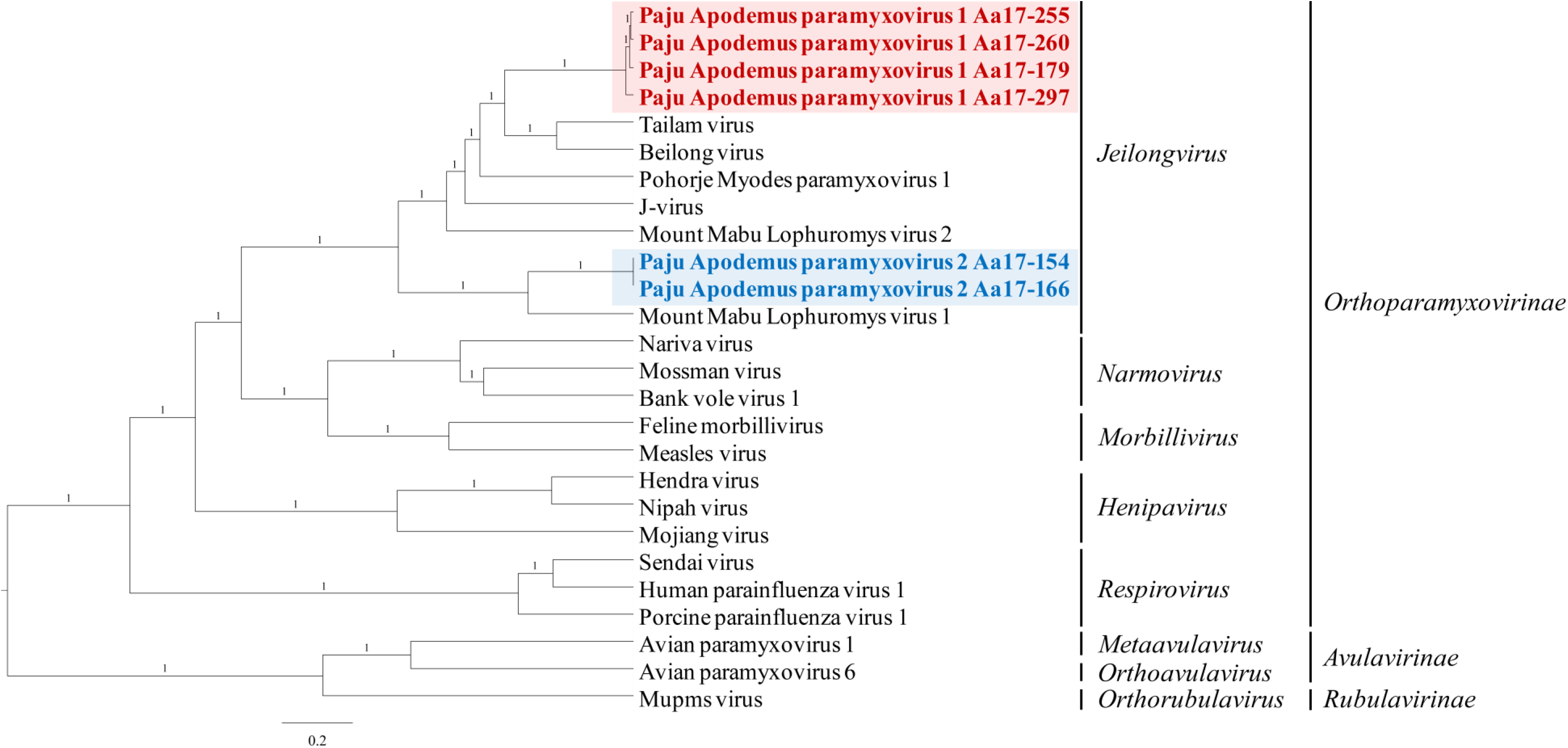
Phylogenetic tree constructed based on the whole-genome sequences of Paju Apodemus paramyxoviruses and other paramyxoviruses. Phylogenetic analysis based on the whole-genome sequences of the Paju Apodemus paramyxoviruses. Evolutionary relationships were inferred using BEAST (v1.10.4) with default priors and assuming homochromous tips. The Markov chain Monte Carlo analysis was performed until adequate sample sizes (ESS >200) were obtained, and TreeAnnotator (v2.5.4) was used to summarize the maximum clade credibility tree from the posterior tree distribution, using a 10% burn-in. *Paramyxoviridae* strains served as reference sequences for the phylogenetic analysis. Red color indicates PAPV-1, and blue indicates PAPV-2.

### Analysis of N-linked glycosylation (NLG) in the G protein of *Jeilongvirus*

To identify the glycosylation patterns of PAPV G proteins, potential NLG sites in the whole amino acid sequences were predicted using NetNglyc 1.0 **(Figure S4)**. PAPV-1 contains 24 NLGs in G proteins; 16/24 NLGs were potentially found at positions 56, 136, 175, 581, 770, 805, 879, 925, 975, 985, 988, 994, 1,052, 1,070, 1,333, and 1,576 over the threshold value (0.5). The ten potential NLG sites from PAPV-2 were estimated, and six of the potential NLGs had significant values at positions 48, 355, 587, 620, 716, and 741. Additionally, the glycosylation pattern of PAPV-1 appeared similar to that of PMPV-1, while the G protein of PAPV-2 was glycosylated less frequently with respect to that of MMLV-1, BeiV, and JV.

### Domain structural analysis of PAPV-1 and -2 G proteins

The primary sequences of PAPV-1 and -2 G proteins are quite different in their lengths. The G protein of PAPV-1 consists of approximately 1,600 amino acids, whereas that of PAPV-2 consists of approximately 700 amino acids. The discrepancy between the two G proteins shows the differential components in the domain diagram **(Figure S5)**. The PAPV-1 G protein contains NH (1–700 aa) and β-strand (1,200–1,600 aa) domains linked by a natively long disordered (DR) region (700–1,200 aa). In contrast, the PAPV-2 G protein has only the NH domain based on protein homology search and secondary structure predictions. Structure prediction and homology modelling indicated that the NH domains in both viruses were largely similar in their protein architecture. Notably, both PAPV-1 and -2 G proteins have a single TM domain in the NH domain. This indicates that the two additional domains in PAPV-1 G protein, DR and β-strand domains, compared to those in PAPV-2 G protein, are most likely expressed as extracellular domains that may interact with host receptor proteins.

### Induction of type I/III IFN, ISGs, and proinflammatory cytokines of PAPV-1 in human epithelial and endothelial cells

PAPV-1 was successfully isolated from Vero E6 cells. To determine the infectivity and induction of antiviral genes in human epithelial and endothelial cells, A549 and HUVEC were infected with PAPV-1, respectively, during 1, 3, 5, and 7 days **(Figures 4 and 5)**. The replication of PAPV-1 gradually increased at 1, 3, 5, and 7 days post-infection(dpi). The mRNA of *Ifnβ*, *ISG15*, *Ifit2/Isg54*, and *Ifit1/Isg56* was upregulated by PAPV-1 infection at 3, 5, and 7 dpi, while *Ifnl1/Il-29* was slightly induced at 7 dpi. The expression of the antiviral genes *Rsad2/Viperin* and *OAS1* increased during PAPV-1 infection, and the cytosolic sensors *Ddx58/Rig-I* and *Ifih1/Mda5* were upregulated at 3, 5, and 7 dpi. In addition, the induction of *Il-6* mRNA was observed in A549 and HUVEC at 3, 5, and 7 dpi. These results demonstrated that PAPV-1 indeed infected human epithelial and endothelial cells and induced the expression of innate antiviral genes including type I/III IFNs, ISGs, and proinflammatory cytokines.

**Figure 4.**
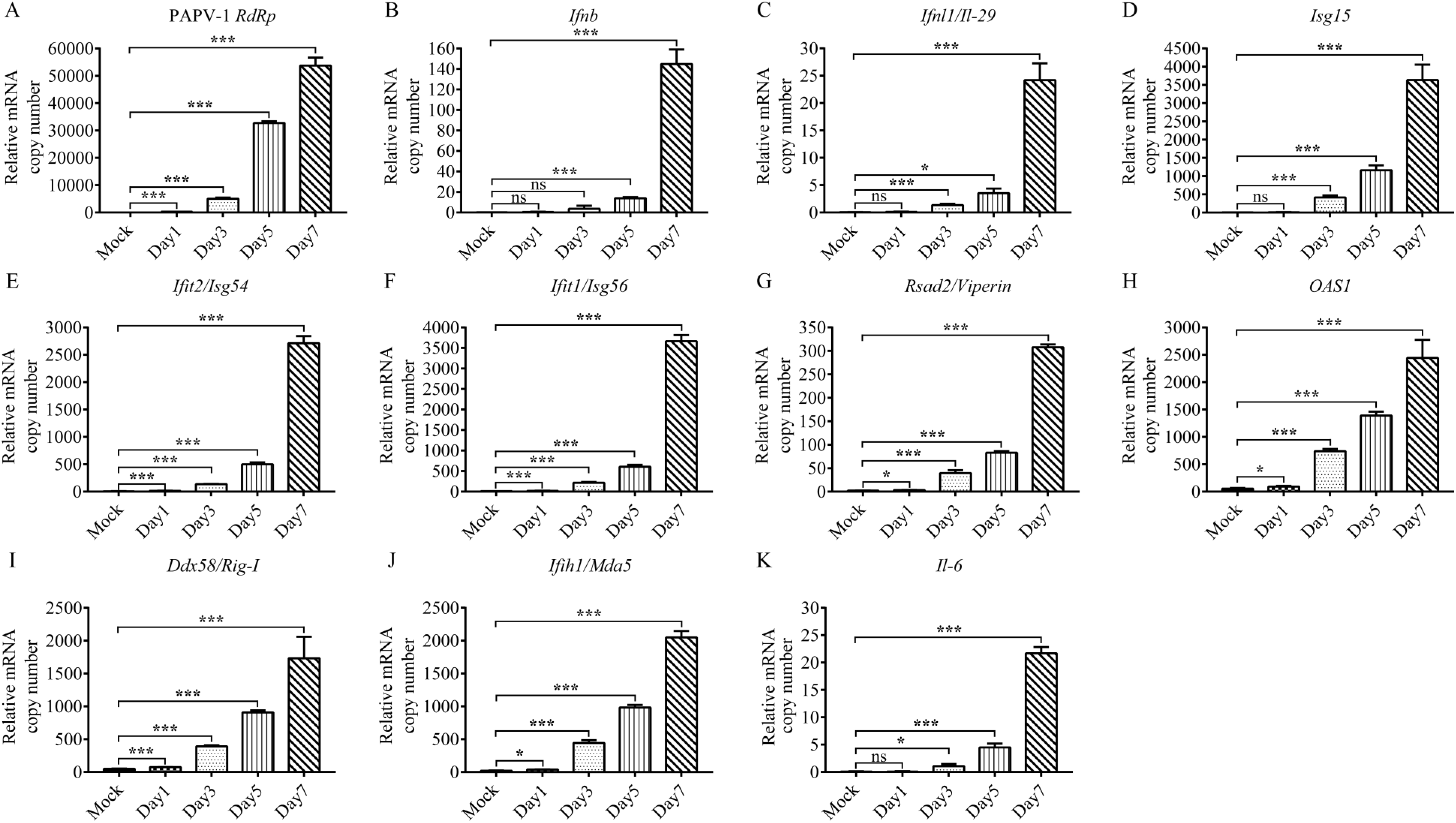
Replication of PAPV-1 and the induction of innate antiviral genes in human epithelial cell (A549) A549 cells were infected with a multiplicity of infection of 0.02 of PAPV-1. Total RNA was analysed via qRT-PCR and examined for the expression of (A) PAPV-1 *RdRp* gene, (B) *Ifnβ*, (C) *Ifnl1/Il-29*, (D) *ISG15*, (E) *Ifit2/Isg54*, (F) *Ifit1/Isg56*, (G) *Rsad2/Viperin*, (H) *OAS1*, (I) *Ddx58/Rig-I*, (J) *Ifih1/Mda5*, and (K) *Il-6* at 1, 3, 5, and 7 days post-infection. Error bars indicate the standard deviation of triplicate measurements in a representative experiment. (*p<0.05; ***p<0.001, unpaired student t-test; ns: non-significant).

**Figure 5.**
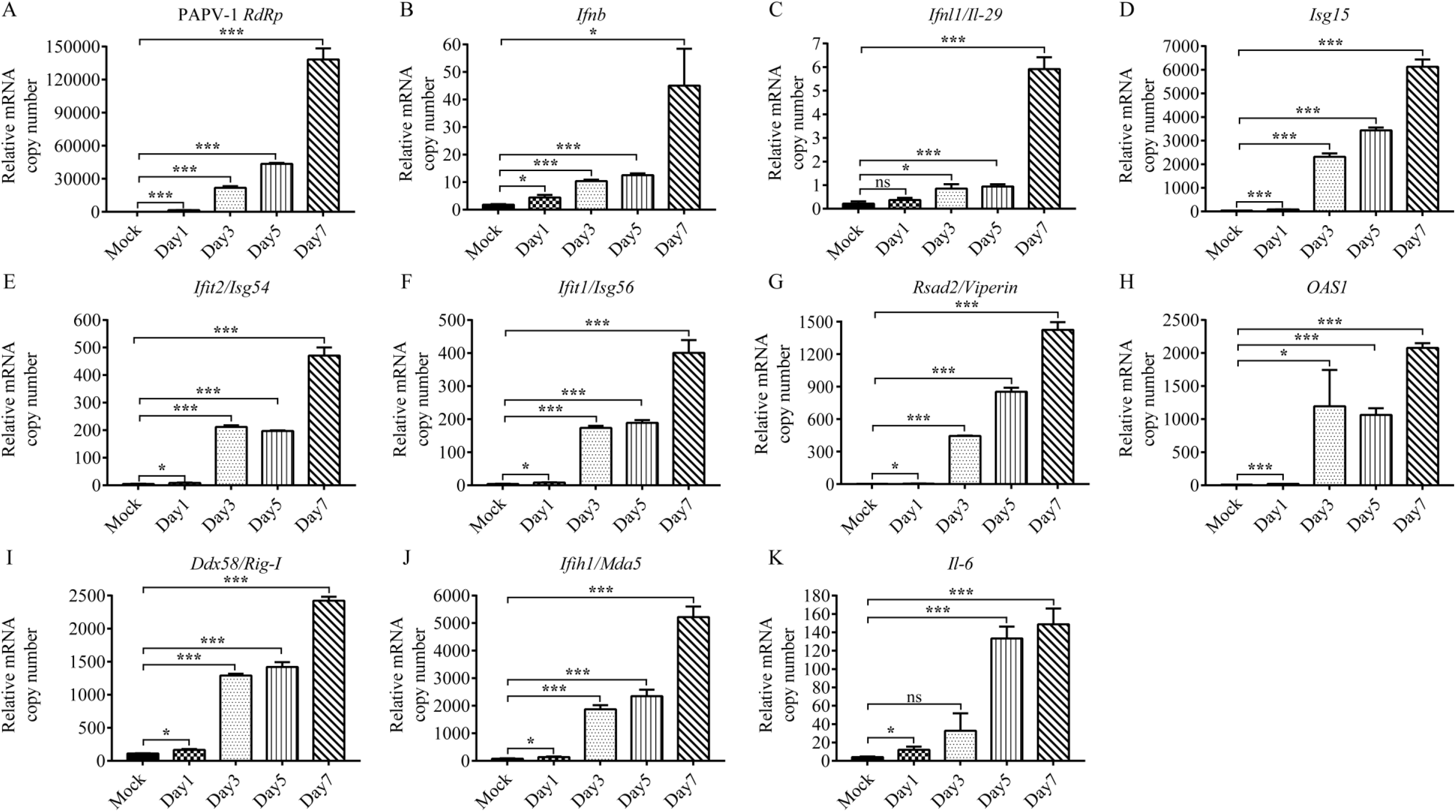
Replication of PAPV-1 and the induction of innate antiviral genes in human endothelial cells (HUVEC) HUVECs were infected with a multiplicity of infection of 0.02 of PAPV-1. Total RNA was analysed via qRT-PCR and examined for the expression of (A) PAPV-1 *RdRp* gene, (C) *Ifnl1/Il-29*, (D) *ISG15*, (E) *Ifit2/Isg54*, (F) *Ifit1/Isg56*, (G) *Rsad2/Viperin*, (H) *OAS1*, (I) *Ddx58/Rig-I*, (J) *Ifih1/Mda5*, and (K) *Il-6* at 1, 3, 5, and 7 days post-infection. Error bars indicate the standard deviation of triplicate measurements in a representative experiment. (*p<0.05; ***p<0.001, unpaired student t-test; ns: non-significant).

## Discussion

Here, we discovered and characterized two novel paramyxoviruses from *A. agrarius* (the striped field mouse) in the ROK. Whole-genome sequences of the paramyxoviruses were obtained using a combination of high-throughput and Sanger sequencing. The phylogenies of PAPV-1 and -2 demonstrated that these two viruses represent distinct genetic lineages within the genus *Jeilongvirus* and family *Paramyxoviridae*. The genome organization (3′-N-P/V/C-M-F-SH-TM-G-L-5′) of these viruses was consistent with that of JV, BeiV, TaiV, and other rodent paramyxoviruses. The genomic characteristics and nucleotide length (19,716 nt) of the PAPV-1 genome are similar to those of PMPV-1, the largest group of rodent-borne paramyxovirues reported to date. This is attributed to the presence of *SH* and *TM* genes and the large size of the *G* gene. Additionally, the complete genome of PAPV-2 was 17,475 nt long, approximately 2 kb shorter than that of PAPV-1, owing to the lack of the *SH* and *TM* genes. According to the paramyxovirus species distinctive criterion (an amino acid distance of >7– 7.5% in the L gene) (25), PAPV-1 and -2 were found to be sufficiently divergent to establish the new genus *Jeilongvirus*. These viruses have been suggested to constitute a separate genus within the family *Paramyxoviridae* (genus *Jeilongvirus*) based on their unique characteristics and evolutionary distance from other paramyxoviruses.

Recently, the host sharing of genetically distinct paramyxoviruses has been reported in nature (18, 26). MMLPV-1 and -2 co-infected in a kidney of a Rungwe brush-furred rat. The bank vole harbored PMPV-1 and bank vole virus (BaVV) in lung and kidney tissues. In this study, PAPV-1 and -2 were first discovered in the same host species, *A. agrarius* although they shared minimal similarity (nucleotide identities of 24.6–24.7%). Notably, paramyxoviruses shared a natural reservoir host with the virus strain belonged to the same family but a significantly distinct phylogenetic lineage, such as PAPV-1 and -2 in *A. agrarius*, MMLPV-1 and -2 in *L. machangui*, and PMPV-1 and BaVV in *M. glareolus*, respectively. These observations arise plausible hypotheses: 1) The virus may be evolved and emerged as a distinctive virus strain via genetic addition or deletion on its progenitor genome. 2) Two naturally distinct viruses might coexist in the same host, followed by competition or cooperation with each other. However, the preferential or predominant emergence of two distinct genotypes of rodent-borne paramyxoviruses awaits further investigation.

Genomic characteristics of paramyxoviruses affect pathogenicity and evolution within hosts (27). Moreover, the molecular prevalence of PAPV-1 was found to be higher than that of PAPV-2. These results led us to hypothesize that the different genome compositions of PAPV-1 and -2 determine their infectivity in nature. First, the SH protein plays a role in viral pathogenicity by affecting the host immune response and membrane fusion mechanism (21–23). The SH protein found in paramyxoviruses, human metapneumovirus (HMPV), and JV modulates TNF-α production and blocks apoptosis *in vitro* and *in vivo*. Deficient SH expression enhances the secretion of proinflammatory cytokines IL-6 and IL-8 compared with that of the wild-type HMPV. The SH protein of HMPV also increases membrane permeability and fusion for viral entry (28). Intriguingly, PAPV-1 was found to possess the *SH* gene, while PAPV-2 showed the absence of the gene. The presence of the *SH* gene may be correlated with the higher prevalence of PAPV-1 (10.6%) in natural hosts compared with that of PAPV-2 (2.6%), since the antagonistic function or increased viral entry promotes propagation in infected cells. Second, the G protein of paramyxoviruses is a predominant determinant of host specificity because it promotes cell entry by interacting with specific proteins on the surface of target cells (29, 30). Different paramyxovirus G proteins have evolved to allow optimal interaction and fusion with target cells in their respective hosts (31–36). The PAPV-1 G protein (1,602 amino acids) was shown to be considerably larger than the PAPV-2 G protein (826 amino acids). The G protein of PAPV-1 consists of the NH, DR, and β-strand domains, while the short length of the PAPV-2 G protein excludes the DR region. Intrinsically, disordered proteins may offer high flexibility to viral proteins either in the wholly or partially disordered form (37). The disordered protein of Zika virus conferred the capability for quick adaption in a changing environment, survival in host body environments, and invasion of the host defence mechanism (38). The characteristics of the DR region of the G protein may influence the higher prevalence of PAPV-1 compared to that of PAPV-2 in nature. Third, the NLG of G protein plays a role in protecting against neutralizing antibodies during cell-cell fusion and viral entry (39). The point mutation of potential NLG sites demonstrated that specific N-glycans in the NiV-G protein are significantly involved in viral entry. NLG in HIV-1 has been associated with survival and immune evasion, such as alteration of sensitivity to neutralizing antibody or reduction of sensitivity to serum antibody (40–42). In the case of NiV, NLG is involved in the proper functioning of proteins and life cycle by having a dual role including enhancement of resistance to antibody neutralization and/or alternative reduction in membrane fusion and viral entry (43). Based on NLG prediction, the G protein of PAPV-1 was found to contain more potential glycosylation sites than that of PAPV-2. Although its precise function is still unclear, the potential glycosylation sites of this protein are thought to aid in shielding the protein from recognition by the host immune system. Thus, the biological consequences of the *SH* gene and molecular characteristics of G protein in PAPV-1 and -2 remain unexplored.

Infectivity and expression of innate antiviral genes significantly influence the pathological effects of viral infection in humans and mice (44–46). Due to the isolation of infectious particles, PAPV-1 was examined for infectivity and induction of innate antiviral genes using human epithelial and endothelial cells. We found that the replication of PAPV-1 increased at 1, 3, 5, and 7 dpi in A549 and HUVEC, respectively. The expression of type I/III IFNs, ISGs, and proinflammatory cytokines were also upregulated in response to PAPV-1. These observations suggest that PAPV-1 may infect and elicit proinflammatory responses in humans. In this study, PAPV-2 was not evaluated owing to the lack of infectious particles. The absence of the PAPV-2 *SH* gene might be involved in the robust induction of antiviral genes including type I IFNs and cytokines, and this might be responsible for the failure to isolate infectious PAPV-2 particles. The comparisons of infectivity, immunogenicity, and pathogenesis between PAPV-1 and -2 remain to be investigated.

In conclusion, we presented two novel paramyxoviruses, PAPV-1 and -2, found in *A. agrarius* in the ROK. These viruses were identified as new Jeilongviruses within the family *Paramyxoviridae* using phylogenetic inference and genomic comparison with the nucleotide and protein sequences of all currently known paramyxovirus species. A total of 102 partial PAPV sequences (83 PAPV-1 and 19 PAPV-2) and six whole genome sequences (four PAPV-1 and two PAPV-2) demonstrated the phylogenetic distribution and relationship of the novel paramyxoviruses in the ROK. PAPV-1 infected human cells and induced the expression of innate antiviral genes. Thus, this study provides profound insights into the molecular prevalence, virus-host interactions, and zoonotic potential of rodent-borne paramyxoviruses. Thus, these observations are expected to raise the awareness of physicians and scientists about the emergence of novel PAPV-1 and -2.

## Materials and Methods

### Ethics statement

The animal trapping procedure was approved by the US Forces Korea (USFK) in accordance with USFK Regulation 40–1 “Prevention, Surveillance, and Treatment of Hemorrhagic Fever with Renal Syndrome.” All procedures and handling of animals were conducted according to the protocol approved by the Korea University Institutional Animal Care and Use Committee (KUIACUC, #2016–0049).

### Animal trapping and PAPVs analyses

Small mammals were captured from 2016 to 2018 using Sherman traps (8 × 9 × 23 cm; H. B. Sherman, Tallahassee, FL, USA). The trapping sites were located in Cheorwon, Chuncheon, Hongcheon, Hwacheon, Inje, Pyeongchang, and Yanggu in Gangwon Province; Dongducheon, Paju, Pocheon, Suwon, Uijeongbu, and Yeoncheon in Gyeonggi Province; Seosan and Taean in Chungcheongnam Province; and Changnyeong in Gyeongsangnam Province. The traps were set at intervals of 1–2 m and examined early the next morning over a period of 1–2 days. Live animals were humanely killed through cardiac puncture under alfaxalone-xylazine anaesthesia and identified to the species level using morphological criteria and PCR when required. A total of 913 rodent species, including 824 *A. agrarius*, 7 *A. peninsulae*, 10 *M. musculus*, 5 *Micromys minutus*, 58 *Myodes regulus*, 9 *Tscherskia triton*, and 158 shrew species were captured. Serum, brain, lung, spleen, kidney, and liver tissues were collected aseptically and frozen at –80°C until use.

### Cell lines

Vero E6 cells (ATCC, #DR-L2785), human lung adenocarcinoma cells (A549) (ATCC, #CCL-185), and human umbilical vein endothelial cells (HUVEC) (ATCC, #CRL1730) were purchased from ATCC. Vero E6, A549, and HUVEC were cultured in DMEM supplemented with Dulbecco’s modified Eagle’s medium (DMEM), 10% fetal bovine serum, 1 mM sodium pyruvate, 2 mL L-glutamine, and 50 mg/ml gentamicin. The cultures were incubated at 37°C in a 5% CO_2_ incubator until use.

### Virus isolation

Kidney tissues were ground in DMEM containing 5% fetal bovine serum. After centrifugation, the supernatant was inoculated into Vero E6 cells. After one and a half hours of adsorption, the excess inoculum was discarded, and the mixture was replaced with 5.5 mL of DMEM. The cultures were incubated at 37°C in a 5% CO_2_ incubator and inspected daily for cytopathic effects using inverted microscopy.

### *In vitro* infection

A total of 1×10^6^ cells per well were prepared in a 6-well plate. After 24 h, the cells were infected with PAPV-1 at a multiplicity of infection of 0.02. The samples were then collected at 1, 3, 5, and 7 days post-infection. Detailed regarding cell lines are available in the supplementary method 4.

### Plaque assay

Vero E6 cells were seeded onto 6-well plates at a density of 1.5×10^6^ cells per well. After overnight incubation at 37°C, the monolayer was washed twice with PBS and inoculated with 10-fold serially diluted viruses. After 90 min of incubation at 37°C with constant shaking, the monolayer was overlaid with a 1:1 overlay medium and medium-melting-point agarose mix. Additionally, following incubation at 37°C for 5 days, the agarose overlay was discarded. The plaques were visualized by staining the monolayer with 0.1% crystal violet in 10% formaldehyde.

### Electron microscopy

Paramyxovirus-infected Vero E6 cells were collected at 7 days post-infection and fixed with 2% paraformaldehyde and 2.5% glutaraldehyde with 0.1 M phosphate buffer, pH 7.4. Thin sections were placed onto 400-mesh square copper electron microscopy grids (Electron Microscopy Sciences) and viewed under a transmission electron microscope (Model H-7650; Hitachi, Japan).

### RNA extraction and RT-PCR

Total RNA was extracted from the lung and kidney tissues of rodents using TRI Reagent Solution (AMBION Inc., Austin, Texas, USA). cDNA was synthesized using a high capacity RNA-to-cDNA kit (Applied Biosystems, Foster City, CA, USA). First, nested PCRs were performed in a 25-μL reaction mixture containing 2.5 U of Ex Taq DNA polymerase (TaKaRa BIO Inc., Shiga, Japan), 2 μg of cDNA, and 10 pM of each primer. The oligonucleotide primer sequences for the nested PCR were PAR-F (outer): 5′-ATG TAY GTB AGT GCW GAT GC-3′, PAR-R1 (outer): 5′-AAC CAD TCW GTY CCR TCA TC-3′, PAR-F and PAR-R2 (inner): 5′-GCR TCR TCW GAR TGR TGD GCA A-3′, and RES-MOR-HEN-F (outer): 5′-TGG GCW GCM AGT GC-3′ and RES-MOR-HEN-R1 (outer): 5′-CCR CAD GCW GTR CAV CCW GT-3′, RES-MOR-HEN-F and RES-MOR-HEN-R2 (inner): 5′-CTG GGT TAC AGC CCC AGC TAC-3′ for the polymerase gene (47). Initial denaturation was performed at 95°C for 5 min, followed by 6 cycles of denaturation at 94℃ for 30 sec, annealing at 37℃ for 30 sec, and elongation at 72℃ for 1 min; followed by 32 cycles of denaturation at 94℃ for 30 sec, annealing at 42℃ for 30 sec, and elongation at 72℃ for 1 min (ProFlex PCR System, Life Technology, CA, USA). PCR products were purified using the LaboPass PCR purification kit (Cosmo Genetech, Seoul, ROK), and sequencing was performed in both directions of each PCR product using a BigDye Terminator v3.1 Cycle Sequencing Kit (Applied Biosystems) on an automated sequencer (ABI 3730XL DNA Analyzer, Applied Biosystems).

### Sequence-independent, single-primer amplification

cDNA was generated from total RNA extracted from paramyxovirus-infected cells using FR26RV-N (5′-GCC GGA GCT CTG CAG ATA TCN NNN NN-3′). The reaction was performed in a 20-μL reaction mixture containing 7 μL total RNA, 2 μL 10 pM of primer, 2 μL 5× First strand buffer, 100 mM dithiothreitol, 25 mM MgCl_2_, 10 mM dNTPs, 0.5 μL RNaseOUT, and 0.5 μL Superscript III RTase (Life Technologies, Carlsbad, CA, USA) in a Proplex thermocycler (Life Technologies). The PCR conditions were as follows: 25°C for 10 min, 50°C for 50 min, and 85°C for 10 min. Double-stranded (ds) cDNA was synthesized using 0.2 units Klenow 3′→5′exo DNA polymerase (Enzynomics, Daejeon, ROK) and 1 μL RNaseH (Invitrogen, San Diego, CA). The Klenow reaction mixture was incubated at 37°C for 1 hr and 75°C for 15 min. The ds cDNA was purified using the MinElute PCR purification kit (Cat No. 28004, Qiagen, Hilden, Germany). Using the FR20RV (5′-GCC GGA GCT CTG CAG ATA TC-3′) primer, ds cDNA was amplified in a 50-μL reaction mixture containing 10 μL ds cDNA template, 10 pM primer, and 2× My Taq Red (Bioline, Taunton, MA, USA). The PCR conditions were as follows: initial denaturation at 98°C for 30 sec, followed by 38 cycles of denaturation at 98°C for 10 sec, annealing at 54°C for 20 sec, and elongation at 72°C for 45 sec.

### NGS for Illumina MiSeq

We prepared libraries using the TruSeq NAano DNA LT Sample Preparation Kit (Illumina, San Diego, CA, USA) according to the manufacturer’s instructions. The samples were mechanically sheared using an M220 focused ultrasonicator (Covaris, Woburn, MA, USA). The cDNA amplicon was size-selected, A-tailed, ligated with indexes and adaptors, and enriched. We sequenced libraries using the MiSeq benchtop sequencer (Illumina) with 2 × 150 bp and a MiSeq reagent V2 (Illumina).

### NGS for Illumina HiSeq

Total RNA was isolated using the Trizol reagent (AMBION). RNA quality was assessed using an Agilent 2100 bioanalyzer (Agilent Technologies, Amstelveen, The Netherlands), and RNA quantification was performed using the ND-2000 Spectrophotometer (Thermo Fisher Scientific, Waltham, MA, USA). Libraries were prepared from total RNA using the NEBNext Ultra II Directional RNA-Seq Kit (NEWENGLAND BioLabs, Ipswich, MA, UK). Additionally, the isolation of mRNA was performed using the Poly(A) RNA Selection Kit (LEXOGEN, Vienna, Austria). The isolated mRNAs were used for cDNA synthesis and shearing, following the manufacturer’s instructions. Indexing was performed using Illumina indexes 1–12. The enrichment step was performed using PCR. Subsequently, the libraries were checked using the Agilent 2100 bioanalyzer (DNA High Sensitivity Kit) to evaluate the mean fragment size. Quantification was performed using a library quantification kit and StepOne Real-Time PCR System (Life Technologies). High-throughput sequencing was performed as paired-end 100 sequencing using the HiSeq X10 system (Illumina).

### NGS data analysis

Adaptor and index sequences of reads were trimmed, and low-quality sequences were filtered using the CLC Genomics Workbench version 7.5.2 (CLC Bio, Cambridge, MA). The genome sequences of TaiV, BeiV, JV, MMLPV-1, and -2 were used in a reference mapping method. Read mapping to the reference genome sequence and extraction of consensus sequences were performed, and the genomic sequences of *Jeilongvirus* were deposited in GenBank (Accession number: MT823459-MT823464). The NGS outputs were analysed using our bioinformatics pipeline. The reads were trimmed with Trimmomatic (v0.36) to remove adapter sequences (48). To exclude the reads from the host genome, they were aligned against the host sequences using Bowtie2 (v2.2.6), and only unaligned reads were used for the subsequent steps (49). Owing to the absence of the completely sequenced genome of the host species, only the complete mitochondrial sequence of the species on the NCBI RefSeq was used as a host reference (50). The remaining reads were filtered for quality using FaQCs (v0.11.5), and de-novo assembly was performed to produce contigs using SPAdes (v3.11.1) (51, 52). The assembled contigs were subsequently examined in a database consisting of complete viral genomes collected from the NCBI RefSeq database (updated in May 2018) using BLASTn (v2.6.0).

### RACE PCR

To obtain the 3′ and 5′ terminal genome sequences of paramyxovirus, we performed RACE PCR using a SMARTer^®^ RACE 5’/3’ Kit (Takara Bio), according to the manufacturer’s specifications. We purified the PCR products using the LaboPass PCR Purification Kit (Cosmo Genetech). Sequencing was performed in both directions of each PCR product using the BigDye Terminator v3.1 Cycle Sequencing Kit (Applied Biosystems) on an automated sequencer (Applied Biosystems).

### Phylogenetic analysis

The viral genomic sequences were aligned and trimmed using the Clustal W tool in the Lasergene program version 5 (DNASTAR, USA), and multiple sequence alignment was performed with high accuracy and high throughput MUSCLE algorithms in MEGA 7.0 (53). Phylogenetic trees were constructed using the maximum likelihood method according to the best-fit substitution model. Support for the topologies was assessed using bootstrapping for 1,000 iterations. In addition, the Bayesian inference method BEAST package (v1.10.4) was used, employing the Markov chain Monte Carlo (MCMC) method (54). The MCMC chain length was set to 100 million states by sampling every 50,000 states. Maximum clade credibility trees were extracted using TreeAnnotator (v1.10.4) and prepared using FigTree (v1.4.0).

### Analysis of potential NLG sites in the G gene

Full-length amino acid sequences were submitted to the NetNlyc 1.0 (Kemitorvet, Denmark) to predict the NLG sites of the G gene of Jeilongviruses (55).

### Domain structural analysis

To find homology, we ran NCBI BLASTP (https://blast.ncbi.nlm.nih.gov/Blast.cgi) using PAPV-1 and -2 G protein sequences against the NR database using default settings. When we ran BLASTP using PAPV-1 G protein and not PAPV-2 G protein, we found the alignments shown in the supplement covering both domains. To detect remote homologs and determine domain architecture, we ran an HHsearch (https://toolkit.tuebingen.mpg.de/tools/hhpred) using PAPV-1 and -2 sequences against PDB, ECOD, and Pfam databases (56, 57). To detect the domain boundaries and various sequence and/or structural features including secondary structure and disordered regions, we ran Quick2D (https://toolkit.tuebingen.mpg.de/tools/quick2d) using PAPV-1 G protein as a query. Finally, to confirm the domain architectures in the related G proteins, we ran Promals with default settings (58).

### Statistical analysis

Statistical analyses were performed as indicated in each figure using GraphPad Prism version 5.00 for Windows (GraphPad Software, San Diego, California, USA; www.graphpad.com).

## Acknowledgments and funding sources

We thank Mr. Su-Am Kim for collecting wild rodents. This work was supported by the Research Program To Solve Social Issues of the National Research Foundation of Korea (NRF) funded by the Ministry of Science and Information and Communication Technology (ICT) (NRF-2017M3A9E4061992 and NRF-2019R1I1A2A01060902). In addition, this work was supported by the Agency for Defense Development (UE202026GD). Partial funding was provided by the Armed Forces Health Surveillance Division Global Emerging Infections Surveillance Branch (GEIS), ProMIS ID P0039_18_ME. The views expressed in this article are those of the author and do not necessarily reflect the official policy or position of the Department of the Army, Department of Defense, or the U.S. Government. Authors, as employees of the U.S. Government (TAK, HCK), conducted the work as part of their official duties. Title 17 U.S.C. §105 provides that ‘Copyright protection under this title is not available for any work of the United States Government.’ Title 17 U.S.C. §101 defines a U.S. Government work is a work prepared by an employee of the U.S. Government as part of the person’s official duties.

## Author Contributions

S.H.L., J.S.N. designed study, collected, analyzed, and interpreted data, and wrote the manuscript. K.K. provided scientific discussion and data analyses. B.H.K., S.C. provided scientific discussion. H.C.K., T.A.K. captured small mammals. S.B., K.P., G.Y.L., H.S.C., S.C., J.W.K., J.G.L., S.H.C. performed experiment and provided scientific discussion and review. C.S.U. provided scientific discussion and review. W.K.K., J.W.S. designed study, analyzed and interpreted data, wrote, reviewed, and revised the manuscript.

