## Supplementary figures for "Novel Paju Apodemus Paramyxovirus 1 and 2, Harbored by *Apodemus agrarius* in The Republic of Korea"

**Supplemental Material**

**Supplemental figure 1. Geographic map of the Republic of Korea showing trapping sites for small mammals from 2016 to 2018**

The map shows the locations of Cheorwon, Chuncheon, Hwacheon, Hongcheon, Inje, and Pyeongchang in Gangwon Province; Dongducheon, Paju, Pocheon, Suwon, Uijeongbu, and Yeoncheon in Gyeonggi Province; Seosan and Taean in Chungcheongnam Province; and Changnyeong in Gyeongsangnam Province. Circles represent each trapping site. Adobe Illustrator CS6 (http://www.adobe.com/products/illustrator.html) was used to construct the map.

**Supplemental figure 2. Phylogenetic trees based on the coding regions (CDS) of the N, P, M, F, G, and L genes of Paju Apodemus paramyxoviruses and other paramyxoviruses**

The amino acid phylogenetic trees of paramyxoviruses were constructed using the maximum likelihood method with distribution models, based on the N, P, M, F, G, and L proteins. Topologies were evaluated using a bootstrap analysis of 1,000 iterations. *Paramyxoviridae* strains served as reference sequences for the phylogenetic analysis. Red color indicates PAPV-1, and blue indicates PAPV-2.

**Supplemental figure 3. Phylogenetic tree based on the partial L sequences of Paju Apodemus paramyxovirus (PAPV)**

Phylogenetic analysis based on 439-bp (coordinates 14,287–14,725 nt) sequences of the Paju Apodemus paramyxoviruses. Evolutionary relationships were inferred using BEAST (v1.10.4) with default priors and assuming homochromous tips. The Markov chain Monte Carlo analysis was performed until adequate sample sizes (ESS >200) were obtained, and TreeAnnotator (v2.5.4) was used to summarize the maximum clade credibility tree from the posterior tree distribution, using a 10% burn-in. *Paramyxoviridae* strains served as reference sequences for the phylogenetic analysis. Red shedding indicates PAPV-1, and blue shedding indicates PAPV-

2. The colors indicate specific sites in the ROK: red, Paju; orange, Yeoncheon; green, Cheorwon; forest green, Pocheon; light blue, Hwacheon; magenta, Yanggu; blue green, Chuncheon; brown, Dongducheon; purple, Seosan; blue, Taean; and pink, Changnyeong.

**Supplemental figure 4. Comparison of consensus-predicted N-linked glycosylation (NLG) sites of the enveloped glycoprotein (G gene) of PAPVs (PAPV-1 and -2) and other representative Jeilongviruses**

To identify glycosylation patterns of G proteins between the PAPVs, potential NLG sites in the whole amino acid sequences of the G gene were predicted using NetNglyc 1.0. The representative Jeilongviruses were Paju Apodemus paramyxovirus 1 MT823459, Beilong virus NC_007803, Tailam virus NC_025355, Mount Mabu Lophuromys paramyxovirus 2 MG573141, Pohorge Myodes paramyxovirus 1 MG516455, J-virus NC_007454, Paju Apodemus paramyxovirus 2 MT823463, and Mount Mabu Lophuromys paramyxovirus 1 MG573140. Red bars represent NLG sites, while black lines indicate thresholds.

**Supplemental figure 5. Structural analysis of Paju Apodemus paramyxoviruses G proteins**

Diagram depicting PAPV-1 and -2 G proteins. Colors indicate domain sites: grey, NH domain; blue, β-strand domain; and red, disordered region

**Supplemental figure 1. Geographic map of the Republic of Korea showing trapping sites for small mammals from 2016 to 2018**

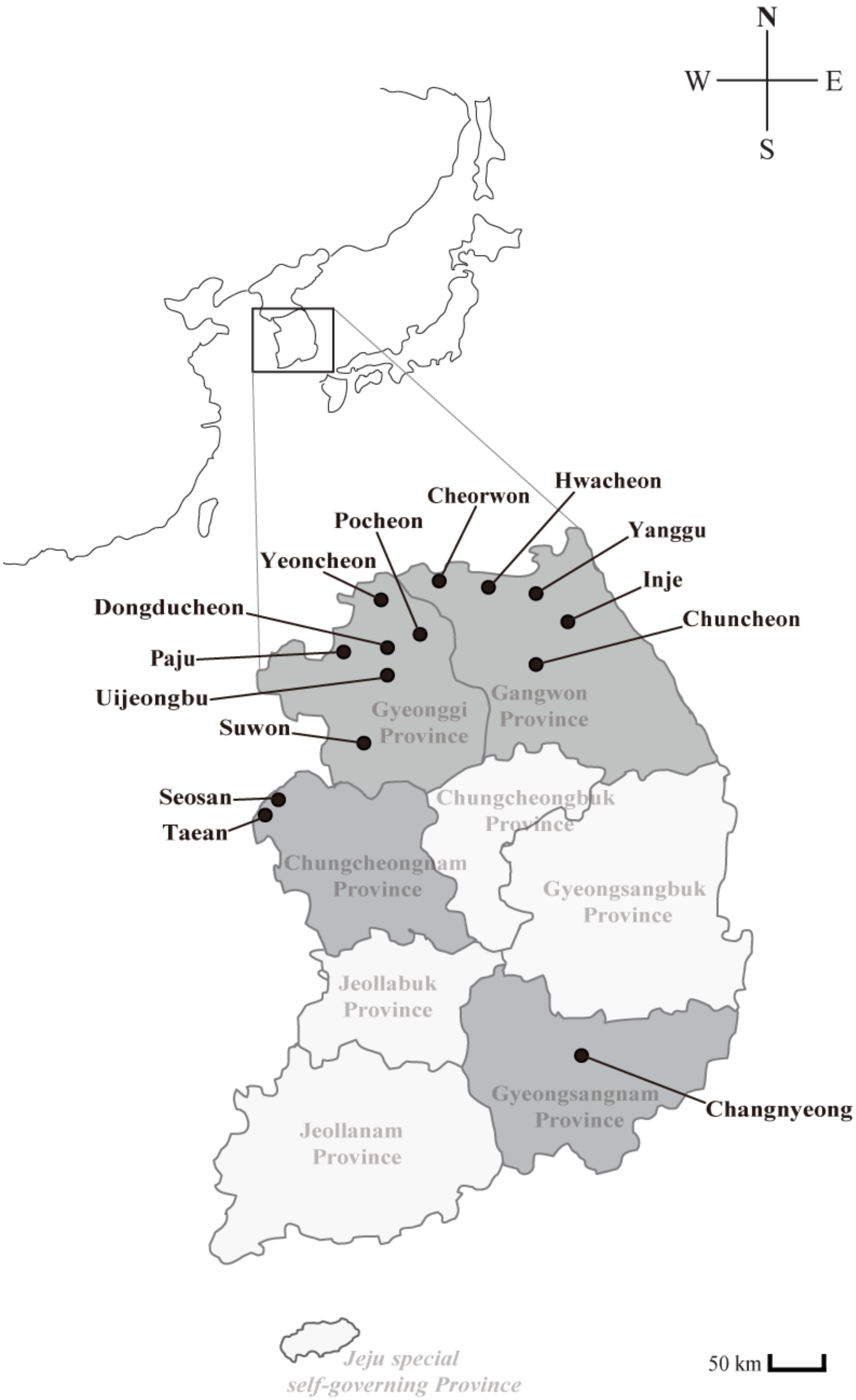

**Supplemental figure 2. Phylogenetic trees based on the coding regions (CDS) of the N, P, M, F, G, and L genes of Paju Apodemus paramyxoviruses and other paramyxoviruses**

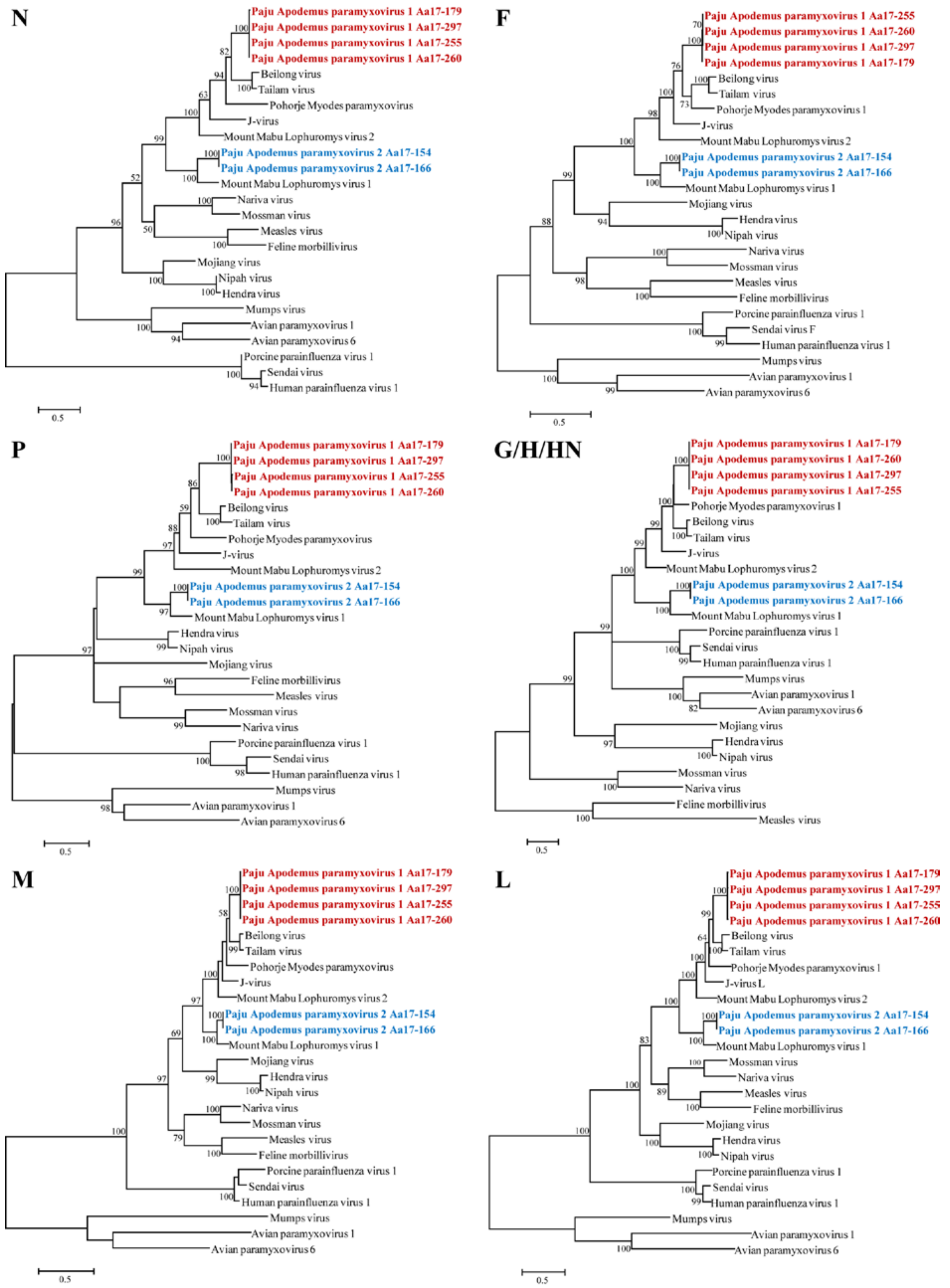

**Supplemental figure 3. Phylogenetic tree based on the partial L sequences of Paju Apodemus paramyxovirus (PAPV)**

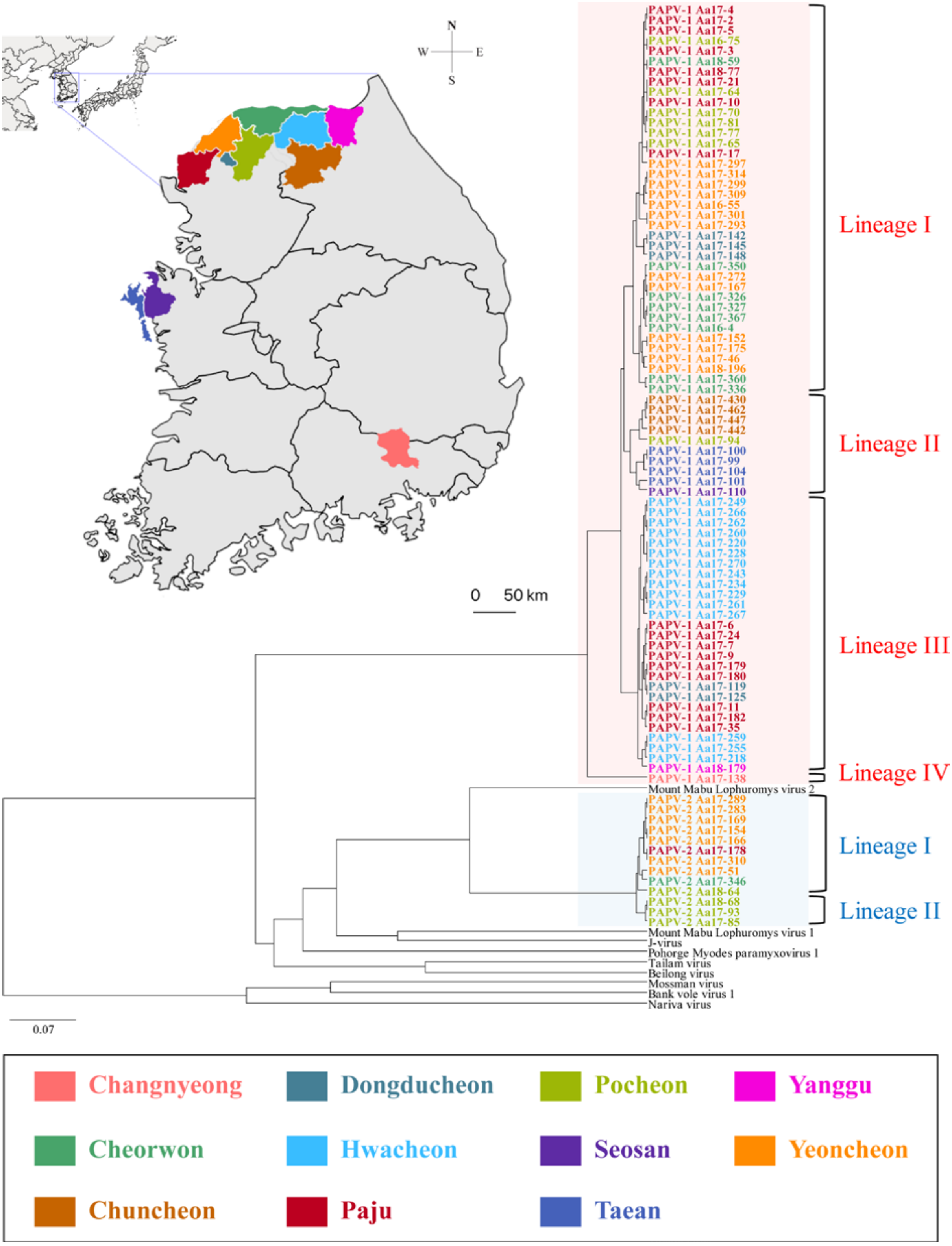

**Supplemental figure 4. Comparison of consensus-predicted N-linked glycosylation (NLG) sites of the enveloped glycoprotein (G gene) of PAPVs (PAPV-1 and -2) and other representative Jeilongviruses**

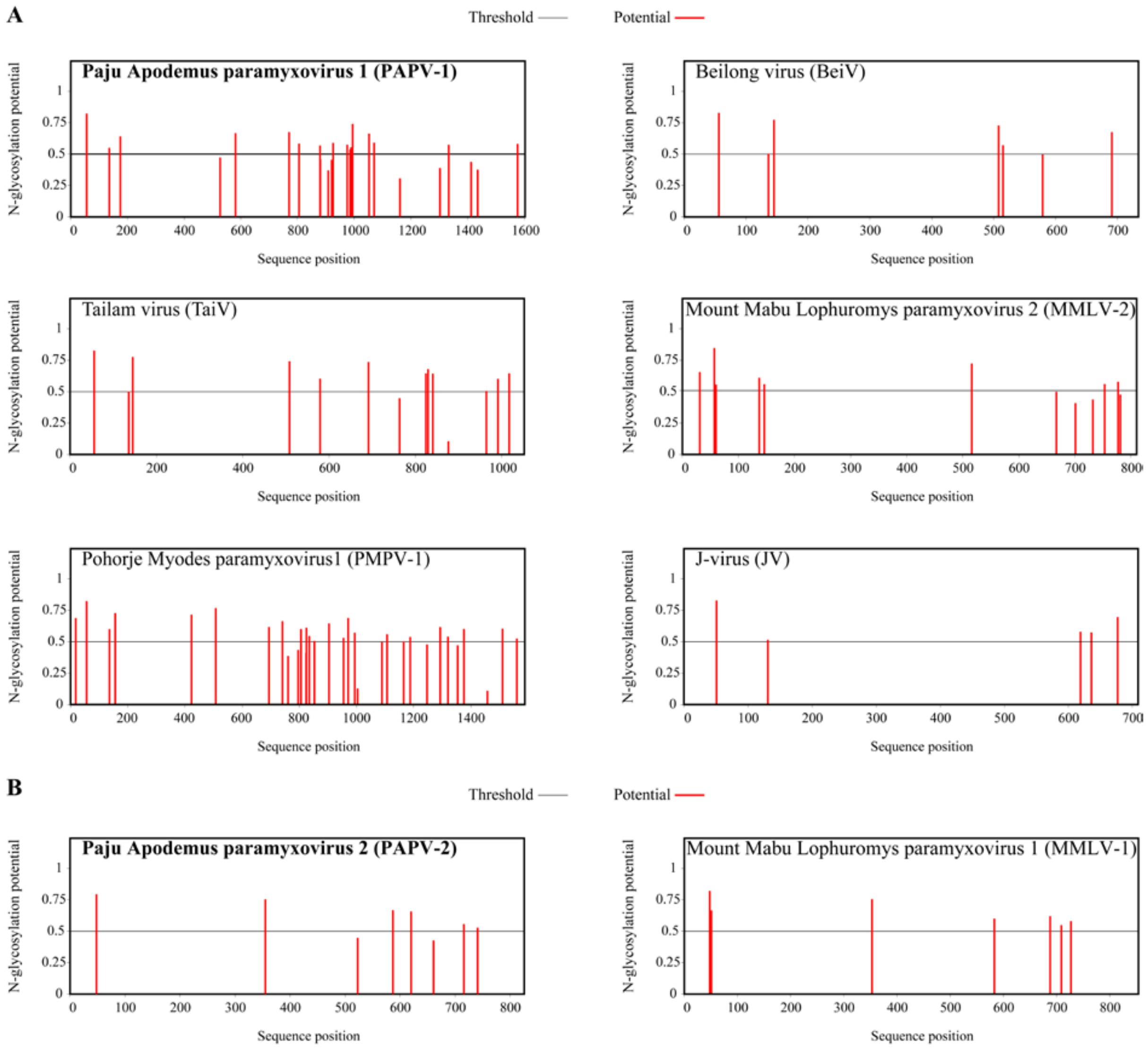

**Supplemental figure 5. Structural analysis of Paju Apodemus paramyxoviruses G proteins**

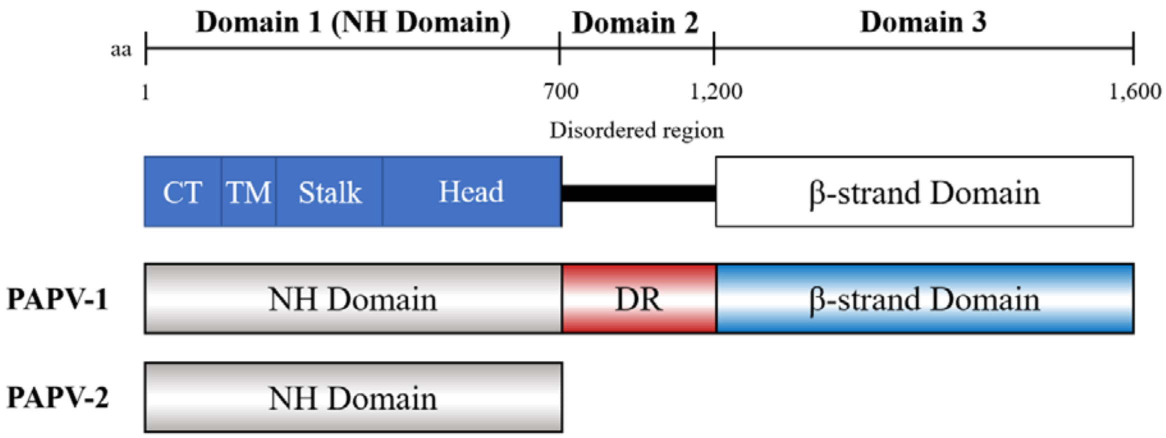
