## Supplementary tables for "Novel Paju Apodemus Paramyxovirus 1 and 2, Harbored by *Apodemus agrarius* in The Republic of Korea"

**Supplemental table 1. Characteristics of Paju Apodemus paramyxovirus (PAPV)-infected *Apodemus agrarius* and the nucleotide sequences of PAPV RNAs acquired in the current study**

| Type of PAPV | Sample | Site (City/Province) | Trapping date | Gender | Weight (g) | Vital status | Nucleotide (nt) position of PAPV RNA |
|---|---|---|---|---|---|---|---|
| PAPV-1 | Aa17-179 [a,b] | Paju/Gyeonggi | Oct. 18, 2017 | Female | 17.0 | Alive | 1–19,716 |
| | Aa17-297 [a,b] | Yeoncheon/Gyeonggi | Nov. 17, 2017 | Female | 15.5 | Alive | 1–19,716 |
| | Aa17-255 [a] | Hwacheon/Gangwon | Nov. 9, 2017 | Male | 26.5 | Dead | 1–19,716 |
| | Aa17-260 [a] | Hwacheon/Gangwon | Nov. 9, 2017 | Female | 17.0 | Alive | 1–19,716 |
| | Aa16-4 | Cheorwon/Gangwon | Jun. 9, 2016 | Female | 5.4 | Alive | 14,287–14,725 |
| | Aa16-14 | Pocheon/Gyeonggi | Jun. 21, 2016 | Female | 25.0 | Alive | 15,369–15,898 |
| | Aa16-55 | Yeoncheon/Gyeonggi | Oct. 12, 2016 | Female | 26.5 | Alive | 14,287–14,725 |
| | Aa16-75 | Pocheon/Gyeonggi | Oct. 19, 2016 | Male | 23.5 | Alive | 14,287–14,725 |
| | Aa16-120 | Hwacheon/Gangwon | Oct. 28, 2016 | Female | 33.5 | Alive | 15,369–15,898 |
| | Aa17-2 | Paju/Gyeonggi | Mar. 29, 2017 | Male | 19.5 | Alive | 14,287–14,725; 15,369–15,898 |
| | Aa17-3 | Paju/Gyeonggi | Mar. 29, 2017 | Male | 22.0 | Dead | 14,287–14,725 |
| | Aa17-4 | Paju/Gyeonggi | Mar. 29, 2017 | Female | 17.5 | Dead | 14,287–14,725 |
| | Aa17-5 | Paju/Gyeonggi | Mar. 29, 2017 | Male | 19.5 | Dead | 14,287–14,725; 15,369–15,898 |
| | Aa17-6 | Paju/Gyeonggi | Mar. 29, 2017 | Female | 16.5 | Alive | 14,287–14,725 |
| | Aa17-7 | Paju/Gyeonggi | Mar. 29, 2017 | Male | 21.5 | Alive | 14,287–14,725; 15,369–15,898 |

| | | | | | | |
|---|---|---|---|---|---|---|
| Aa17-9 | Paju/Gyeonggi | Mar. 29, 2017 | Female | 20.5 | Alive | 14,287–14,725 |
| Aa17-10 | Paju/Gyeonggi | Mar. 29, 2017 | Male | 20.5 | Alive | 14,287–14,725 |
| Aa17-11 | Paju/Gyeonggi | Mar. 29, 2017 | Male | 38.5 | Alive | 14,287–14,725 |
| Aa17-17 | Paju/Gyeonggi | Mar. 29, 2017 | Male | 27.0 | Alive | 14,287–14,725; 15,369–15,898 |
| Aa17-21 | Paju/Gyeonggi | Mar. 29, 2017 | Male | 29.5 | Alive | 14,287–14,725; 15,369–15,898 |
| Aa17-24 | Paju/Gyeonggi | Mar. 29, 2017 | Male | 23.0 | Alive | 14,287–14,725; 15,369–15,898 |
| Aa17-35 | Paju/Gyeonggi | Mar. 29, 2017 | Female | 16.0 | Alive | 14,287–14,725 |
| Aa17-46 | Yeoncheon/Gyeonggi | Apr. 18, 2017 | Female | 25.5 | Alive | 14,287–14,725 |
| Aa17-64 | Pocheon/Gyeonggi | Apr. 27, 2017 | Male | 25.0 | Alive | 14,287–14,725 |
| Aa17-65 | Pocheon/Gyeonggi | Apr. 27, 2017 | Male | 27.0 | Alive | 14,287–14,725; 15,369–15,898 |
| Aa17-70 | Pocheon/Gyeonggi | Apr. 27, 2017 | Male | 27.5 | Alive | 14,287–14,725; 15,369–15,898 |
| Aa17-77 | Pocheon/Gyeonggi | Apr. 27, 2017 | Female | 27.0 | Dead | 14,287–14,725; 15,369–15,898 |
| Aa17-81 | Pocheon/Gyeonggi | Apr. 27, 2017 | Female | 22.5 | Dead | 14,287–14,725; 15,369–15,898 |
| Aa17-94 | Pocheon/Gyeonggi | Apr. 28, 2017 | Female | 22.5 | Dead | 14,287–14,725 |
| Aa17-99 | Taean/Chungcheongnam | May 17, 2017 | Male | 21.5 | Alive | 14,287–14,725; 15,369–15,898 |
| Aa17-100 | Taean/Chungcheongnam | May 17, 2017 | Female | 31.0 | Alive | 14,287–14,725 |
| Aa17-101 | Taean/Chungcheongnam | May 17, 2017 | Male | 28.5 | Alive | 14,287–14,725; 15,369–15,898 |

| | | | | | | |
|---|---|---|---|---|---|---|
| Aa17-104 | Taean/Chungcheongnam | May 17, 2017 | Female | 31.5 | Alive | 14,287–14,725; 15,369–15,898 |
| Aa17-110 | Seosan/Chungcheongnam | May 18, 2017 | Male | 30.5 | Alive | 14,287–14,725; 15,369–15,898 |
| Aa17-119 | Dongducheon/Gyeonggi | May 24, 2017 | Female | 29.5 | Alive | 14,287–14,725; 15,369–15,898 |
| Aa17-125 | Dongducheon/Gyeonggi | May 25, 2017 | Male | 30.5 | Alive | 14,287–14,725; 15,369–15,898 |
| Aa17-138 | Changnyeong/Gyeongsangnam | Jun. 25, 2017 | Female | 30.5 | Alive | 14,287–14,725 |
| Aa17-142 | Dongducheon/Gyeonggi | Sep. 13, 2017 | Female | 34.0 | Alive | 14,287–14,725 |
| Aa17-145 | Dongducheon/Gyeonggi | Sep. 13, 2017 | Male | 38.0 | Alive | 14,287–14,725 |
| Aa17-148 | Dongducheon/Gyeonggi | Sep. 14, 2017 | Male | 31.5 | Alive | 14,287–14,725 |
| Aa17-152 | Yeoncheon/Gyeonggi | Sep. 28, 2017 | Male | 42.0 | Alive | 14,287–14,725 |
| Aa17-167 | Yeoncheon/Gyeonggi | Sep. 28, 2017 | Male | 37.0 | Alive | 14,287–14,725 |
| Aa17-175 | Yeoncheon/Gyeonggi | Sep. 29, 2017 | Male | 40.0 | Alive | 14,287–14,725 |
| Aa17-180 | Paju/Gyeonggi | Oct. 18, 2017 | Male | 26.0 | Alive | 14,287–14,725 |
| Aa17-182 | Paju/Gyeonggi | Oct. 18, 2017 | Female | 36.0 | Alive | 14,287–14,725 |
| Aa17-203 | Paju/Gyeonggi | Oct. 27, 2017 | Male | 34.5 | Dead | 14,287–14,725 |
| Aa17-207 | Paju/Gyeonggi | Oct. 27, 2017 | Male | 30.5 | Alive | 15,369–15,898 |
| Aa17-218 | Hwacheon/Gangwon | Nov. 8, 2017 | Male | 30.5 | Alive | 15,369–15,898 |
| Aa17-220 | Hwacheon/Gangwon | Nov. 8, 2017 | Male | 28.0 | Alive | 14,287–14,725; 15,369–15,898 |

| | | | | | | |
|---|---|---|---|---|---|---|
| Aa17-228 | Hwacheon/Gangwon | Nov. 8, 2017 | Male | 19.5 | Alive | 14,287–14,725 |
| Aa17-229 | Hwacheon/Gangwon | Nov. 8, 2017 | Male | 15.5 | Alive | 14,287–14,725; 15,369–15,898 |
| Aa17-234 | Hwacheon/Gangwon | Nov. 8, 2017 | Male | 31.5 | Alive | 14,287–14,725; 15,369–15,898 |
| Aa17-243 | Hwacheon/Gangwon | Nov. 8, 2017 | Male | 16.0 | Alive | 14,287–14,725; 15,369–15,898 |
| Aa17-249 | Hwacheon/Gangwon | Nov. 9, 2017 | Male | 18.5 | Dead | 14,287–14,725; 15,369–15,898 |
| Aa17-259 | Hwacheon/Gangwon | Nov. 9, 2017 | Female | 17.0 | Dead | 14,287–14,725; 15,369–15,898 |
| Aa17-261 | Hwacheon/Gangwon | Nov. 9, 2017 | Female | 19.0 | Alive | 14,287–14,725; 15,369–15,898 |
| Aa17-262 | Hwacheon/Gangwon | Nov. 9, 2017 | Male | 20.0 | Alive | 14,287–14,725 |
| Aa17-266 | Hwacheon/Gangwon | Nov. 9, 2017 | Female | 30.0 | Alive | 14,287–14,725 |
| Aa17-267 | Hwacheon/Gangwon | Nov. 9, 2017 | Male | 15.0 | Alive | 14,287–14,725 |
| Aa17-270 | Hwacheon/Gangwon | Nov. 9, 2017 | Male | 34.0 | Alive | 14,287–14,725 |
| Aa17-272 | Yeoncheon/Gyeonggi | Nov. 16, 2017 | Female | 23.5 | Dead | 14,287–14,725; 15,369–15,898 |
| Aa17-293 | Yeoncheon/Gyeonggi | Nov. 17, 2017 | Female | 40.0 | Alive | 14,287–14,725 |
| Aa17-299 | Yeoncheon/Gyeonggi | Nov. 17, 2017 | Male | 23.0 | Alive | 14,287–14,725 |
| Aa17-301 | Yeoncheon/Gyeonggi | Nov. 17, 2017 | Female | 25.5 | Dead | 14,287–14,725; 15,369–15,898 |
| Aa17-309 | Yeoncheon/Gyeonggi | Nov. 17, 2017 | Female | 27.0 | Dead | 14,258–14,697 |
| Aa17-313 | Yeoncheon/Gyeonggi | Nov. 17, 2017 | Male | 28.0 | Dead | 15,369–15,898 |

| | | | | | | |
|---|---|---|---|---|---|---|
| Aa17-314 | Yeoncheon/Gyeonggi | Nov. 17, 2017 | Male | 20.0 | Dead | 14,287–14,725 |
| Aa17-326 | Cheorwon/Gangwon | Apr. 6, 2017 | Female | 22.0 | Dead | 14,287–14,725; 15,369–15,898 |
| Aa17-327 | Cheorwon/Gangwon | Apr. 6, 2017 | Male | 26.5 | Dead | 14,287–14,725; 15,369–15,898 |
| Aa17-336 | Cheorwon/Gangwon | Apr. 6, 2017 | Male | 29.0 | Dead | 14,287–14,725; 15,369–15,898 |
| Aa17-345 | Cheorwon/Gangwon | Apr. 17, 2017 | Male | 15.5 | Dead | 14,287–14,725; 15,369–15,898 |
| Aa17-350 | Cheorwon/Gangwon | Oct. 11, 2017 | Male | 39.0 | Dead | 14,287–14,725 |
| Aa17-360 | Cheorwon/Gangwon | Oct. 23, 2017 | Female | 33.5 | Dead | 14,287–14,725 |
| Aa17-367 | Cheorwon/Gangwon | Oct. 23, 2017 | Male | 35.5 | Dead | 14,287–14,725; 15,369–15,898 |
| Aa17-388 | Chuncheon/Gangwon | Oct. 16, 2017 | Female | 30.2 | Dead | 15,369–15,898 |
| Aa17-409 | Chuncheon/Gangwon | Oct. 25, 2017 | Female | 43.1 | Dead | 15,369–15,898 |
| Aa17-430 | Chuncheon/Gangwon | Oct. 30, 2017 | Female | 20.4 | Dead | 15,369–15,898 |
| Aa17-442 | Chuncheon/Gangwon | Oct. 31, 2017 | Female | 23.1 | Dead | 14,287–14,725; 15,369–15,898 |
| Aa17-447 | Chuncheon/Gangwon | Oct. 31, 2017 | Male | 38.0 | Dead | 14,287–14,725 |
| Aa17-456 | Chuncheon/Gangwon | Nov. 8, 2017 | Male | 16.6 | Dead | 14,287–14,725 |
| Aa17-461 | Chuncheon/Gangwon | Nov. 12, 2017 | Female | 24.8 | Dead | 14,287–14,725 |
| Aa17-462 | Chuncheon/Gangwon | Nov. 12, 2017 | Female | 23.5 | Dead | 14,287–14,725 |
| Aa18-59 | Cheorwon/Gangwon | Sep. 11, 2018 | Male | 38.0 | Alive | 14,287–14,725; 15,369–15,898 |

| | | | | | | | |
|---|---|---|---|---|---|---|---|
| | Aa18-77 | Paju/Gyeonggi | Oct. 16, 2018 | Female | 35.0 | Dead | 14,287–14,725 |
| | Aa18-179 | Yanggu/Gangwon | Nov. 28, 2018 | Female | 26.0 | Alive | 14,287–14,725 |
| | Aa18-183 | Yanggu/Gangwon | Nov. 28, 2018 | Female | 14.0 | Alive | 15,369–15,898 |
| | Aa18-196 | Yeoncheon/Gyeonggi | Dec. 4, 2018 | Male | 25.5 | Alive | 14,287–14,725 |
| | Aa17-154 [a] | Yeoncheon/Gyeonggi | Sep. 28, 2017 | Female | 37.5 | Alive | 1–17,476 |
| | Aa17-166 [a] | Yeoncheon/Gyeonggi | Sep. 28, 2017 | Male | 36.5 | Alive | 1–17,476 |
| | Aa16-32 | Yeoncheon/Gyeonggi | Oct. 11, 2016 | Female | 44.5 | Dead | 15,369–15,898 |
| | Aa17-49 | Yeoncheon/Gyeonggi | Apr. 18, 2017 | Male | 26.0 | Dead | 15,369–15,898 |
| | Aa17-51 | Yeoncheon/Gyeonggi | Apr. 18, 2017 | Female | 28.0 | Alive | 14,287–14,725 |
| | Aa17-85 | Pocheon/Gyeonggi | Apr. 28, 2017 | Female | 25.5 | Alive | 14,287–14,725; 15,369–15,898 |
| PAPV-2 | Aa17-93 | Pocheon/Gyeonggi | Apr. 28, 2017 | Female | 26.0 | Dead | 14,287–14,725 |
| | Aa17-169 | Yeoncheon/Gyeonggi | Sep. 29, 2017 | Female | 34.5 | Dead | 14,287–14,725 |
| | Aa17-178 | Paju/Gyeonggi | Oct. 18, 2017 | Female | 47.5 | Alive | 14,287–14,725 |
| | Aa17-283 | Yeoncheon/Gyeonggi | Nov. 16, 2017 | Female | 32.0 | Dead | 14,287–14,725; 15,369–15,898 |
| | Aa17-289 | Yeoncheon/Gyeonggi | Nov. 16, 2017 | Female | 42.0 | Dead | 14,287–14,725; 15,369–15,898 |
| | Aa17-300 | Yeoncheon/Gyeonggi | Nov. 16, 2017 | Male | 6.0 | Dead | 15,369–15,898 |
| | Aa17-310 | Yeoncheon/Gyeonggi | Nov. 17, 2017 | Male | 32.5 | Dead | 14,287–14,725 |

| | | | | | | |
|---|---|---|---|---|---|---|
| Aa17-335 | Cheorwon/Gangwon | Apr. 6, 2017 | Female | 24.0 | Dead | 15,369–15,898 |
| Aa17-346 | Cheorwon/Gangwon | Apr. 17, 2017 | Male | 28.5 | Dead | 14,287–14,725; 15,369–15,898 |
| Aa17-348 | Cheorwon/Gangwon | Apr. 17, 2017 | Male | 21.5 | Dead | 15,369–15,898 |
| Aa17-390 | Chuncheon/Gangwon | Oct. 17, 2017 | Female | 16.1 | Dead | 15,369–15,898 |
| Aa18-64 | Pocheon/Gyeonggi | Sep. 12, 2018 | Male | 35.0 | Alive | 14,287–14,725 |
| Aa18-68 | Pocheon/Gyeonggi | Sep. 12, 2018 | Male | 44.5 | Dead | 14,287–14,725 |
| Aa18-78 | Paju/Gyeonggi | Oct. 16, 2018 | Female | 39.5 | Dead | 15,369–15,898 |
| Aa18-84 | Paju/Gyeonggi | Oct. 16, 2018 | Female | 32.0 | Alive | 15,369–15,898 |

[a]: Whole genome sequences were completely obtained.
[b]: Cell culture-based isolates.

**Supplemental table 2. Molecular prevalence of Paju Apodemus paramyxoviruses (PAPVs) by region, sex, weight, and season in the Republic of Korea from 2016 to 2018**

| Category | Number of *A. agrarius* | RNA positivity (%)[*] | | |
|---|---|---|---|---|
| | | PAPV-1 | PAPV-2 | Total |
| **Region (n=824)** | | | | |
| Gangwon | 361 | 35/361 (9.7%) | 4/361 (1.1%) | **39/361 (10.8%)** |
| Gyeonggi | 434 | 46/434 (10.6%) | 17/434 (3.9%) | **63/434 (14.5%)** |
| Gyeongsangnam | 6 | 1/6 (16.7%) | 0/6 | **1/6 (16.7%)** |
| Chungcheongnam | 23 | 5/23 (21.7%) | 0/23 | **5/23 (21.7%)** |
| **Sex (n=824)** | | | | |
| Males | 370 | 48/370 (13.0%) | 8/370 (2.2%) | **56/370 (15.1%)** |
| Females | 453 | 39/453 (8.6%) | 13/453 (2.9%) | **52/453 (11.5%)** |
| Unknown | 1 | 0/1 | 0/1 | **0/1** |
| **Weight (n=824)**[**] | | | | |
| <10 g | 22 | 0/22 | 1/22 (4.5%) | **1/22 (4.5%)** |
| ≤10–20 g | 344 | 22/344 (6.4%) | 1/344 (0.3%) | **23/344 (6.7%)** |
| ≤20–30 g | 248 | 38/248 (15.3%) | 7/248 (2.8%) | **45/248 (18.1%)** |
| ≤30–40 g | 152 | 26/152 (17.1%) | 8/152 (5.3%) | **34/152 (22.4%)** |
| ≤40 g | 58 | 1/58 (1.7%) | 4/58 (6.9%) | **5/58 (8.6%)** |
| **Season (n=824)** | | | | |
| Spring (Mar.–May) | 208 | 31/208 (14.9%) | 7/208 (3.4%) | **38/208 (18.3%)** |
| Summer (Jun.–Aug.) | 37 | 3/37 (8.1%) | 0/37 | **3/37 (8.1%)** |
| Autumn (Sep.–Nov.) | 564 | 52/564 (9.2%) | 14/564 (2.5%) | **66/564 (11.7%)** |
| Winter (Dec.–Feb.) | 15 | 1/15 (6.7%) | 0/15 (%) | **1/15 (6.7%)** |
| **Total** | **824** | **87/824 (10.6%)** | **21/824 (2.6%)** | **108/824 (13.1%)** |

[*]The positive rate of PAPV RNA indicates the detection of the partial L gene (targeting pan-*Orthoparamyxovirinae* and the genera *Respirovirus*, *Morbillivirus*, and *Henipavirus*) using RT-PCR and Sanger sequencing.

[**]Statistically significant (Chi-square exact test, p<0.05)

**Supplemental table 3. Genome coverage of consensus sequences for Paju Apodemus paramyxoviruses (PAPVs) using next generation sequencing (NGS) methods**

| NGS method | Origin | Strains | Total reads | Read mapped to reference/Total reads (%) | PAPVs | |
|---|---|---|---|---|---|---|
| | | | | | Read mapped to reference | Depth of coverage* |
| SISPA-based MiSeq | Vp | Aa17-179 [a] | 1,623,052 | 8.892 | 144,317 | 142230.86 |
| | | Aa17-297 [a] | 1,479,714 | 5.362 | 79,344 | 78394.25 |
| **Total** | | | **3,102,766** | **7.208** | **223,661** | **111830.5** |
| RNA-Seq based HiSeq | Kidney | Aa17-255 [a] | 89,469,566 | 0.004 | 3,859 | 3848.04 |
| | | Aa17-260 [a] | 89,095,350 | 0.027 | 23,807 | 23767.15 |
| | | Aa17-154 [b] | 83,021,016 | 0.004 | 3,584 | 3576.00 |
| | | Aa17-166 [b] | 106,752,990 | 0.049 | 52,367 | 52355.01 |
| **Total** | | | **368,338,922** | **0.021** | **20,904** | **20886.55** |

*Depth of coverage was calculated using the number of mapped reads (read length×number of reads that matches the reference/reference genome size).

Vp: Cell culture-based isolates

[a]: Paju Apodemus paramyxovirus 1

[b]: Paju Apodemus paramyxovirus 2

SISPA: sequence-independent, single-primer amplification

**Supplemental table 4. Sequences of intergenic regions (IGR) and transcriptional start and stop signals of Paju Apodemus paramyxoviruses (PAPVs)**

| **Genes** | **Gene stop** | **IGR** | **Gene Start** |
|---|---|---|---|
| /N | | CTT | AGGAGCAAAG |
| N/P | TTAAGAAAAA | CTT | AGGAGTCAAG |
| P/M | TTAAGAAAAA | CTT | AGGAGGAAGG |
| M/F | CTAAGAAAAA | CTT | AGGATTCAAA |
| F/SH* | TTACGAAAAA | CTT | AGGACCAAAG |
| SH/TM | TAATAAAAAA | CTT | AGGGCAAATG |
| TM/G | TTAAGAAAAA | CTT | AGGACGAAAG |
| G/L | TTAAGAAAAA | CTT | AGGATGAATG |
| L/ | TTAAGAAAAA | CTT | |
| Consensus sequences | YWAHRAAAAA | CTT | AGGRBNMADR |

*PAPV-1 has the *SH* gene, while PAPV-2 does not.

**Supplemental table 5. Amino acid similarities between Paju Apodemus paramyxoviruses (PAPVs) and other species of *Paramyxoviridae***

| Genus | Virus | PAPV-1 | | | | | | | | PAPV-2 | | | | | | |
|---|---|---|---|---|---|---|---|---|---|---|---|---|---|---|---|---|
| | | N | P | M | F | SH[a] | TM[a] | G | L | N | P | M | F | TM[a] | G | L |
| *Jeilongvirus* | **PAPV-1** | 99.6 | 99.2 | 100 | 99.6 | 97.8 | 99.2 | 97.6 | 99.5 | 43.9 | 32.2-32.9 | 72.0-72.2 | 56.8-56.9 | 14.3 | 28.5-30.6 | 61.8 |
| | **PAPV-2** | 43.9 | 32.2-32.9 | 72.0-72.2 | 56.8-56.9 | - | 14.3 | 28.5-30.6 | 61.8 | 100 | 100 | 100 | 100 | 99.5 | 99.9 | 100 |
| | TaiV | 62.8 | 51.2 | 83.5 | 72.2-72.3 | 32.5 | 34.1 | 46.1-51.5 | 77.7-77.8 | 42.3 | 32.0 | 71.7-71.9 | 53.7-53.8 | 18.0 | 29.6 | 61.6 |
| | BeiV | 61.9 | 52.4 | 83.2 | 72.6 | 32.5 | 31.4 | 60.8-60.9 | 77.3 | 42.3 | 31.4 | 72.0-72.2 | 52.4-52.5 | 18.9 | 30.1 | 60.8 |
| | JV | 53.3-53.4 | 46.8-47.0 | 80.3 | 71 | 12.9-14.3 | 26.5-27.2 | 48.2-48.4 | 74.0 | 43.7 | 32.0 | 71.4-71.6 | 53.9-54.0 | 16.6 | 31.5 | 61.1-61.2 |
| | MMLV-1 | 42.8-44.5 | 30.1-31.6 | 70 | 53.3 | - | 13.7 | 28.9-30.0 | 61.3 | 60.7-62.3 | 60.5 | 86.1-86.4 | 73.0-73.1 | 42.4 | 45.2 | 79.7-79.8 |
| | MMLV-2 | 54.0-54.2 | 35.7-35.9 | 77.9 | 60.7-60.8 | - | 28.4 | 40.1-41.6 | 70.0-70.1 | 42.6 | 30.3-30.4 | 71.7-71.9 | 54.2-54.3 | 17.5 | 29.9 | 61.8 |
| | PMPV-1 | 54.0-54.2 | 45.8 | 77.1 | 70.0 | 18.1-19.3 | 29.2 | 35.2-51.8 | 73.4 | 42.9-43.2 | 34.0-34.1 | 67.8-68.0 | 56.1-56.2 | 11.5 | 30.7 | 61.0 |
| *Henipavirus* | HeV | 33.7 | 24.0-24.8 | 50.6 | 37.2-37.4 | - | - | 19.0-19.2 | 47.3-47.5 | 35.7 | 24.0-24.1 | 51.6-51.8 | 33.1 | - | 17.5 | 47.5 |
| | NiV | 33.5 | 22.0-23.2 | 50.9 | 39.0-39.2 | - | - | 17.9-18.1 | 7.2-47.4 | 37.0 | 23.4-23.5 | 52.8-23.0 | 34.2-34.3 | - | 18.1 | 47.5 |
| | MojV | 35.2 | 22.2-23.6 | 53.2 | 37.2 | - | - | 50.5 | 48.8-48.9 | 32.1 | 22.6-22.7 | 55.8-55.9 | 38.8-38.9 | - | 16.2 | 50.0 |
| *Morbillivirsu* | FeMV | 33.7 | 19.1-19.3 | 47.8 | 33.7-33.9 | - | - | 12.1 | 46.1 | 32.9 | 18.3-18.4 | 49.0 | 32.4 | - | 12.4 | 47.5 |
| | MV | 34.7 | 17.1 | 45.1 | 32.1 | - | - | 9.6-1.5 | 47.0 | 33.7 | 20.2 | 45.1 | 32.5-32.6 | - | 11.0 | 47.7-47.8 |
| *Narmovirus* | MossV | 35.6 | 20.2-21.0 | 47.9 | 31.5 | - | - | 13.3-13.6 | 50.4-50.5 | 36.4 | 21.8 | 49.9-50.0 | 30.1-30.2 | - | 13.6 | 51.3-51.4 |
| | NarV | 35.6 | 20.2-20.4 | 49.1 | 33.3-33.7 | - | - | 13.1-14.9 | 48.5 | 38.7 | 22.8-22.9 | 52.2-52.4 | 30.7-30.8 | - | 14.0 | 50.2-50.3 |
| *Respirovirus* | PPIV-1 | 24.0-24.3 | 11.3-13.3 | 36.8 | 27.5 | - | - | 24.1-24.3 | 37.8-38.0 | 20.9-21.1 | 12.8-12.9 | 35.7-35.8 | 26.7 | - | 24.8 | 37.9 |
| | SenV | 20.2 | 14.9-15.5 | 34.4 | 28.1 | - | - | 23.8-24.0 | 37.9 | 21.0 | 14.1 | 37.5-37.6 | 26.8-26.9 | - | 27.0 | 38.1-38.2 |
| | HPIV-1 | 20.6 | 12.1-12.7 | 37.1 | 26.8 | - | - | 23.8-24.0 | 39.2-39.5 | 22.5 | 9.6 | 38.9-39.1 | 28.1-28.2 | - | 25.6 | 38.6 |
| *Orthorubulavirus* | MuV | 22.0-23.7 | 14.3 | 17.6 | 25.3-25.7 | - | - | 21.3 | 29.0-29.1 | 24.2-24.7 | 11.6 | 18.6 | 24.0 | - | 23.4 | 29.7 |
| *Orthoavulavirus* | APMV-1 | 25.4 | 12.2-15.2 | 17.1 | 23.5 | - | - | 20.3 | 26.0 | 25.2 | 11.6 | 19.8 | 25.0 | - | 19.8 | 26.6 |
| *Metaavulavirus* | APMV-6 | 30.3 | 12.6-12.8 | 19.4 | 28.4 | - | - | 17.0 | 27.3-27.4 | 28.0 | 16.5 | 20.6-20.7 | 28.3-28.4 | - | 18.3 | 27.8 |

Similarities are indicated per protein, expressed as percentages. HeV, Hendra virus; NiV, Nipah virus; MojV, Mòjiāng virus; FeMV, Feline morbillivirus; MeV, Measles virus; PPIV-1, Porcine parainfluenza virus 1; SenV, Sendai virus; HPIV-1, Human parainfluenza virus 1; MuV, Mumps virus; APMV-1, Avian paramyxovirus 1; APMV-6, Avian paramyxovirus 6

[a]Not present in all viruses

**Supplemental table 6. Sequences of primers for RT-qPCR**

| Genes | Forward (Sense, 5'–3') | Reverse (Antisense, 5'–3') |
|---|---|---|
| *Human b-actin* | AGAGCTACGAGCTGCCTGAC | CGTGGATGCCACAGGACT |
| *Human GAPDH* | GCAAATTCCATGGCACCGT | TCGCCCCACTTGATTTTGG |
| *Ifnβ* | GTCAGAGTGGAAATCCTAAG | ACAGCATCTGCTGGTTGAAG |
| *Il-29* | CGCCTTGGAAGAGTCACTCA | GAAGCCTCAGGTCCCAATTC |
| *Isg15* | GCGAACTCATCTTTGCCAGTA | CCAGCATCTTCACCGTCAG |
| *Ifit2/Isg54* | ATCCCCCATCGCTTATCTCT | CCACCTCAATTAATCAGGCACT |
| *Ifit1/Isg56* | GGATTCTGTACAATACACTAGAAACCA | CTTTTGGTTACTTTTCCCCTATCC |
| *Rsad2/Viperin* | TGCTTTTGCTTAAGGAAGCTG | TCTACTTTGCAGAACCTCACCA |
| *OAS-1* | GAAGGCAGCTCACGAAACC | AGGCCTCAGCCTCTTGTG |
| *Ddx58/Rig-I* | CAGACAGATCCGAGACACTA | TGCAAGACCTTTGGCCAGTT |
| *Ifih1/Mda5* | GAGGAATCAGCACGAGGAATAA | TCAGATGGTGGGCTTTGAC |
| *Il-6* | GCCCAGCTATGAACTCCTTCT | GCGGCTACATCTTTGGAATCT |
| PAPV-1 *RdRp* | GGCCGCCTTGTATATTTCC | GGCCTCATTATTTGAATGTGC |
